# Characterizing the reproductive transcriptomic correlates of acute dehydration in males in the desert-adapted rodent, *Peromyscus eremicus*

**DOI:** 10.1101/096057

**Authors:** Lauren Kordonowy, Matthew MacManes

**Author notes:** **Corresponding Author:**Lauren Kordonowy, Department of Molecular, Cellular, and Biomedical Sciences, University of New Hampshire, Rudman Hall (MCBS), 46 College Road, Durham, NH, 03824, (802)735-5849.

## Abstract

The understanding of genomic and physiological mechanisms related to how organisms living in extreme environments survive and reproduce is an outstanding question facing evolutionary and organismal biologists. One interesting example of adaptation is related to the survival of mammals in deserts, where extreme water limitation is common. Research on desert rodent adaptations has focused predominantly on adaptations related to surviving dehydration, while potential reproductive physiology adaptations for acute and chronic dehydration have been relatively neglected. This study aims to explore the reproductive consequences of acute dehydration by utilizing RNAseq data in the desert-specialized cactus mouse (*Peromyscus eremicus*). Specifically, we exposed 22 male cactus mice to either acute dehydration or control (fully hydrated) treatment conditions, quasimapped testes-derived reads to a cactus mouse testes transcriptome, and then evaluated patterns of differential transcript and gene expression. Following statistical evaluation with multiple analytical pipelines, nine genes were consistently differentially expressed between the hydrated and dehydrated mice. We hypothesized that male cactus mice would exhibit minimal reproductive responses to dehydration; therefore, this low number of differentially expressed genes between treatments aligns with current perceptions of this species’ extreme desert specialization. However, these differentially expressed genes include Insulin-like 3 (Insl3), a regulator of male fertility and testes descent, as well as the solute carriers Slc45a3 and Slc38a5, which are membrane transport proteins that may facilitate osmoregulation. Together, these results suggest that in male cactus mice, acute dehydration may be linked to reproductive modulation via Insl3, but not through gene expression differences in the subset of other *a priori* tested reproductive hormones. Although water availability is a reproductive cue in desert-rodents exposed to chronic drought, potential reproductive modification via Insl3 in response to acute water-limitation is a result which is unexpected in an animal capable of surviving and successfully reproducing year-round without available external water sources. Indeed, this work highlights the critical need for integrative research that examines every facet of organismal adaptation, particularly in light of global climate change, which is predicted, amongst other things, to increase climate variability, thereby exposing desert animals more frequently to the acute drought conditions explored here.

## Background

For decades, evolutionary biologists have successfully described examples where natural selection has resulted in the exquisite match between organism and environment (*e.g*. Salinity adaptations in three-spine sticklebacks: Hohenlohe et al., 2010; Jones et al. 2012; high-altitude adaptations for hemoglobin in deer mice and humans: Storz et al., 2010, Lorenzo et al., 2015; and *Peromyscus* adaptations for multiple environments: Hoekstra et al., 2006; Bedford & Hoekstra, 2015; Munshi-South & Richardson, 2016). The match between organism and environment must be studied in the context of both components of fitness: survival and reproductive success, because both aspects of selection are critical to long term persistence in a given environment. Habitat specialists must possess phenotypes enabling survival and successful reproduction; therefore, cases where environmental selective pressures result in reduced reproductive success (*e.g*. Martin & Wiebe, 2004; Bolger, Patten & Bostock, 2005; Evans et al., 2010; Wingfield, Kelley & Angelier, 2011), but not survival, demand attention. Species occupying extreme environments are likely more vulnerable to the bifurcation of these two components of fitness. Moreover, long-term events like global climate change are predicted to increase climate variability and may enhance the challenges faced by species living on the fringes of habitable environments (Martin & Wiebe, 2004; Somero, 2010; Wingfield, Kelley & Angelier, 2011; Wingfield, 2013; Asres & Amha, 2014).

Deserts present extraordinary environmental impediments for habitation, including extreme heat, aridity, and solar radiation. Examples of well-described desert mammal behavioral adaptations are seasonal torpor (reviewed in Kalabukhov 1960; Geiser, 2010), nocturnality (e.g. Stephens & Tello, 2009; Fuller et al., 2014) and burrowing (reviewed in Vorhies, 1945; Kelt, 2011) to avoid high temperatures and sun exposure. Desert mammals also exhibit a wide range of morphological adaptations, including large ears for effective heat dissipation (e.g. Schmidt-Nieslen, 1964; Hill & Veghte, 1976), metabolic water production (e.g. MacMillen & Hinds, 1983; reviewed in Walsberg 2000), and renal adaptations to minimize water-loss (e.g. Schmidt-Nielsen et al., 1948; Dantzler, 1982; Diaz, Ojeda & Rezenda, 2006). Although desert rodents must possess adaptations conferring survival *and* reproductive benefits, researchers have focused on their physiological adaptations for survival. For example, renal adaptations in species of Kangaroo rats (*Dipodomys* species) have been described and explored for over 60 years (Schmidt-Nielsen et al., 1948; Schmidt-Nielsen and Schmidt-Nielsen, 1952; Marra et al., 2012; Urity et al., 2012). While early research determined the renal physiology for Kangaroo rats (Schmidt-Nielsen et al., 1948; Schmidt-Nielsen and Schmidt-Nielsen, 1952; Vimtrup and Schmidt-Nielsen), recent research has focused on the genetic underpinnings of this phenotype (Marra et al., 2012; Urity et al., 2012; Marra, Romero & DeWoody, 2014; Marra et al., 2014), which is indicative of a larger methodological shift in the approach for examining adaptation.

Research in another desert-adapted rodent, *Peromyscus eremicus* (cactus mouse), has followed a somewhat different trajectory; however, it too has only pursued survival oriented physiological mechanisms (but see Kordonowy and MacManes, 2016; Kordonowy et al., 2017; MacManes, 2017). The ecology, physiology and behaviors of the cactus mouse in comparison with other *Peromyscus* species were summarized in 1968 (King, ed.), and the relationships between basal metabolic rate, body mass, and evaporative water loss were reviewed several decades later (MacMillen and Garland, 1989). Known desert adaptations for cactus mouse include nocturnality and torpor (reviewed in Veal and Caire, 1979; Caire, 1999); however, the cactus mouse does not possess the same elaborate kidney structures responsible for renal adaptations in kangaroo rats (Dewey, Elias & Appel, 1966; MacManes 2016, *unpublished data*). The physiological renal adaptations in *P. eremicus* have not been described in detail, despite considerable explorations of other aspects of this species’ biology (reviewed in Veal and Caire, 1979; Caire, 1999). In order to initially characterize renal function of the cactus mouse, water consumption measurements and electrophysical dehydration effects for this species have also recently been documented (Kordonowy et al., 2017). Because the renal mechanisms for mitigating renal water-loss in *P. eremicus* have not been determined, a comparative genetic approach may be instrumental for characterizing this species’ adaptive kidney phenotype. To this end, MacManes and Eisen (2014) conducted a comparative analysis to find genes expressed in the kidney tissue of cactus mouse that were under positive selection relative to other mammals. MacManes (2017) also recently conducted differential gene expression analyses on cactus mouse kidneys subjected to acute dehydration to explore transcriptomic renal responses. However, the transcriptomic resources available for this species extend considerably beyond renal tissue; transcripts from cactus mouse (as well as numerous other *Peromyscus* species) have been heavily utilized to pursue questions related to multiple aspects of evolutionary biology (reviewed in Bedford and Hoekstra, 2015; Munshi-South and Richardson, 2016). Current investigations into cactus mouse desert-adaptive renal physiology include transcriptomic analyses (MacManes 2017); however, we extended this genetic approach by shifting the focus from adaptions for survival to include physiological adaptations for reproductive success (Kordonowy and MacManes, 2016). The cactus mouse is an ideal system for investigating dehydration effects on reproduction, as well as potential reproductive adaptations for drought, given decades of study of reproductive biology, as well as more recent development of transcriptomic resources that include male reproductive tissues.

Substantial research has been done on the effects of various types of stress on reproduction (e.g. Wingfield & Sapolsky, 2003; Ahmed et al., 2015; Nargund, 2015; Wingfield, 2013); furthermore, the impacts of dehydration stress on reproduction in desert specialized rodents have been historically explored by studies documenting the impacts of water availability as a reproductive cue (reviewed in Schwimmer and Heim 2009; Bales and Hostetler, 2011). Specifically, some female desert rodents have shown evidence of reproductive attenuation due to water-limitation (Mongolian gerbil: Yahr and Kessler, 1975; hopping mouse: Breed, 1975), and male Mongolian gerbils subject to dehydration had decreased reproductive tissue mass (Yahr and Kessler, 1975). In contrast, Shaw’s jird, an Egyptian desert rodent, did not elicit perceivable reproductive response to water deprivation in either males or females (El, Bakry et al., 1999). Furthermore, water-supplementation studies among wild desert rodents resulted in prolonged breeding seasons in the hairy-footed gerbil and the four-striped grass mouse, but not in the Cape short-eared gerbil (Christian, 1979). Recent research has confirmed the importance of rainfall as a reproductive cue in the Arabian spiny mouse (Sarli et al., 2016), the Baluchistan gerbil (Sarli et al., 2015), Chessman’s gerbil (Henry and Dubost, 2012) and the Spinifex hopping mouse (Breed and Leigh, 2011). The focus of this previous research was to investigate reproductive cues and consequences of water-limitation in desert rodents, namely how species have adapted breeding onset and cessation patterns to respond to water availability. Our current study experimentally tests reproductive responses to acute dehydration using a differential gene expression approach in the cactus mouse, which has not been previously evaluated for reproductive impacts of dehydration.

In nature, wild cactus mice are subjected to both acute and chronic dehydration, and understanding the reproductive effects of dehydration stress is an initial step for fully characterizing the suite of phenotypes enabling their successful reproduction. Given that this species has evolved in southwestern United States deserts and it breeds continuously throughout the year (Veal and Caire, 1979; Caire, 1999), we predict that neither acute nor chronic water stress, while physiologically demanding, would be associated with reproductive suppression. To evaluate acute water stress reproductive tissue gene expression responses in the current study, we leveraged previous research that characterized the transcriptome of male *P. eremicus* reproductive tissues from functional and comparative perspectives (Kordonowy and MacManes, 2016). We extend upon this work by performing an RNAseq experiment to identify differentially expressed genes in testes between male *P. eremicus* subjected to acute dehydration versus control (fully hydrated) animals in order to determine the impacts, if any, on male reproduction. We hypothesized that male cactus mice would exhibit minimal gene expression level reproductive responses to acute dehydration because they are highly desert-adapted and they breed year-round, including in times of chronic draught. Specifically, we predicted that genes linked to reproductive function would not be differentially expressed in the testes in response to acute dehydration. We pursued this line of research on the effects of dehydration on reproduction in cactus mouse in order to begin to address the need for additional studies focusing on physiological adaptations related to reproductive success in animals living in extreme, and changing, environments.

## Methods

### Treatment Groups, Sample Preparation and mRNA Sequencing

The cactus mice used for this study include only captive born individuals purchased from the *Peromyscus* Genetic Stock Center (Columbia, South Carolina). The animals at the stock center are descendant from individuals originally collected from a hot-desert location in Arizona more than 30 years ago. The colony used in this study has been housed since 2013 at the University of New Hampshire in conditions that mimic temperature and humidity levels in southwestern US deserts, as described previously (Kordonowy & MacManes, 2016). Males and females are housed together, which provides olfactory cues to support reproductive maturation. Males do not undergo seasonal testicular atrophy, as indicated by successful reproduction throughout the year. The individuals used in this study were all of the same developmental stage – reproductively mature – which was assessed by observing that the testes had descended into the scrotum from the abdomen, making them visible.

Males that had free access to water prior to euthanasia are labeled as WET mice in our analyses. Mice that were water deprived, which we refer to as DRY mice, were weighed and then water deprived for ∼72 hours directly prior to euthanasia. All mice were weighed prior to sacrifice, and DRY mice were evaluated for weight loss during dehydration. Individuals in the study were collected between September 2014 – April 2016.

Cactus mice were sacrificed via isoflurane overdose and decapitation in accordance with University of New Hampshire Animal Care and Use Committee guidelines (protocol number 130902) and guidelines established by the American Society of Mammalogists (Sikes et al., 2016). Trunk blood samples were collected following decapitation for serum electrolyte analyses with an Abaxis Vetscan VS2 using critical care cartridges (Abaxis). The complete methodology and results of the electrolyte study, as well as the reported measures of water consumption and weight loss due to dehydration are described fully elsewhere (Kordonowy et al., 2016). Rather, this study focused on differential gene expression between the testes of 11 WET and 11 DRY mice. Testes were harvested within ten minutes of euthanasia, placed in RNAlater (Ambion Life Technologies), flash-frozen in liquid nitrogen, and stored at -80° degree Celsius. A TRIzol, chloroform protocol was implemented for RNA extraction (Ambion Life Technologies). Finally, the quantity and quality of the RNA product was evaluated with both a Qubit 2.0 Fluorometer (Invitrogen) and a Tapestation 2200 (Agilent Technologies, Palo Alto, USA).

Libraries were made with a TruSeq Stranded mRNA Sample Prep LT Kit (Illumina), and the quality and quantity of the resultant sequencing libraries were confirmed with the Qubit and Tapestation. Each sample was ligated with a unique adapter for identification in multiplex single lane sequencing. We submitted the multiplexed samples of the libraries for processing on lanes at the New York Genome Center Sequencing Facility (NY, New York). Paired end sequencing reads of length 125bp were generated on an Illumina 2500 platform. Reads were parsed by individual samples according to their unique hexamer IDs in preparation for analysis.

### Assembly of Testes Transcriptome

We assembled a testes transcriptome from a single reproductively mature male using the *de novo* transcriptome protocol described previously (MacManes, 2016). The testes transcripts were assembled with alternative methodologies utilizing several optimization procedures to produce a high-quality transcriptome; however, the permutations of this assembly process are described extensively elsewhere (MacManes, 2016; Kordonowy and MacManes, 2016). The testes transcriptome we selected was constructed as described below. The raw reads were error corrected using Rcorrector version 1.0.1 (Song & Florea, 2015), then subjected to quality trimming (using a threshold of PHRED <2, as per MacManes 2014) and adapter removal using Skewer version 0.1.127 (Jiang et al, 2014). These reads were then assembled in the *de novo* transcriptome assembler BinPacker version 1.0 (Liu et al., 2016). We also reduced sequence redundancy to improve the assembly using the sequence clustering software CD-HIT-EST version 4.6 (Li & Godzik, 2006; Fu et al., 2012). We further optimized the assembly with Transrate version 1.0.1 (Smith-Unna et al., 2015) by retaining only highly supported contigs (cutoff: 0.02847). We then evaluated the assembly’s structural integrity with Transrate and assessed completeness using the vertebrata database in BUSCO version 1.1b 1 (Simão et al., 2015). We quasimapped the raw reads to the assembly with Salmon version 0.7.2 (Patro, Duggal & Kingsford, 2015) to confirm that mapping rates were high. Finally, the assembly was also annotated in dammit version 0.3.2, which finds open reading frames with TransDecoder and uses five databases (Rfam, Pfam, OrthoDB, BUSCO, and Uniref90) to thoroughly annotate transcripts (https://github.com/camillescott/dammit).

### Differential Gene and Transcript Expression Analyses

Several recent studies have critically evaluated alternative methodologies for differential transcript and gene expression to determine the relative merits of these approaches (Gierlinski et al., 2015; Schurch et al., 2016; Soneson, Love & Robinson, 2016; Froussios et al., 2016). Soneson and colleagues (2016) demonstrated that differential gene expression (DGE) analyses produce more accurate results than differential transcript expression (DTE) analyses. Furthermore, the differential gene expression approach is more appropriate than differential transcript expression for the scope of our research question, which is true of many evolutionary genomic studies (Soneson et al., 2016). However, because both DTE and DGE approaches are widespread in current literature, we deemed it important to confirm that these methodologies yielded concordant results in the current study.

We utilized edgeR (Robinson, McCarthy & Smith, 2010; McCarthy, Chen & Smith, 2012) as our primary statistical software because Schurch and colleagues (2016) rigorously tested various packages for analyzing DGE, and edgeR performed optimally within our sample size range. While edgeR is a widely used statistical package for evaluating differential expression, we also confirmed our results with another popular package, DESeq2 (Love, Huber & Anders, 2014), in order to validate our findings.

We performed differential expression analyses with three alternative methodologies. Two analyses were conducted in R version 3.3.1 (R Core Team, 2016) using edgeR version 3.16.1, a Bioconductor package (release 3.4) that evaluates statistical differences in count data between treatment groups (Robinson, McCarthy & Smith, 2010; McCarthy, Chen & Smith, 2012). Our first method utilized tximport, an R package developed by Soneson and colleagues (2016), which incorporates transcriptome mapping-rate estimates with a gene count matrix to enable downstream DGE analysis. The authors assert that such transcriptome mapping can generate more accurate estimates of DGE than traditional pipelines (Soneson et al, 2016). While our first methodology evaluated differential gene expression, our second analysis used the transcriptome mapped read sets to perform differential transcript expression and identify the corresponding gene matches. The purpose of this second analysis was to evaluate whether the transcript expression results coincided with the gene expression results produced by the same program, edgeR. Finally, our third methodology determined differential gene expression with tximport in conjunction with DESeq2 version 1.14.0 (Love, Huber & Anders, 2014), a Bioconductor package (release 3.4) which also evaluates statistical differences in expression. We performed this alternative DGE analysis with DESeq2 in order to corroborate our DGE results from edgeR. Thus, the results for all three differential expression analyses were evaluated to determine the coincidence among the genes identified as significantly different between the WET and DRY groups. These alternative differential expression methods are described in detail below.

We quasimapped each of the 11 WET and 11 DRY sample read sets to the testes transcriptome with Salmon version 0.7.2 to generate transcript count data. To perform the gene-level analysis in edgeR, we constructed a gene ID to transcript ID mapping file, which was generated by a BLASTn (Altschul et al., 1990; Madden, 2002) search for matches in the *Mus musculus* transcriptome (ensembl.org) version 7/11/16 release-85. We then imported the Salmon-generated count data and the gene ID to transcript ID mapping file into R using the tximport package (Soneson et al. 2016) to convert the transcript count data into gene counts. This gene count data was imported into edgeR for differential gene expression analysis (Robinson, McCarthy & Smith, 2010; McCarthy, Chen & Smith, 2012). We applied TMM normalization to the data, calculated common and tagwise dispersions, and performed exact tests (p < 0.05) adjusting for multiple comparisons with the Benjamini-Hochburg correction (Benjamini & Hochburg, 1995) to find differentially expressed genes, which we identified in Ensembl (ensemble.org).

Next, we performed a transcript-level analysis using edgeR. To accomplish this, the Salmon-generated count data was imported into R and analyzed as was described above for the gene-level analysis in edgeR. After determining which transcript IDs were differentially expressed, we identified the corresponding genes using the gene ID to transcript ID matrix described previously. The significantly expressed transcripts without corresponding gene matches were selected for an additional BLASTn search in the NCBI non-redundant nucleotide database (http://blast.ncbi.nlm.nih.gov/Blast.cgi). However, these results were not subjected to any additional analyses, because these matches were not consistent across all three differential expression analyses. This list of BLASTn search matches is provided in supplementary materials (DTEno-matchBLASTnSequences.md).

The third analysis used DESeq2 to conduct an additional gene-level test, using the same methods as described for the previous gene-level analysis, with the exception that data were imported into an alternative software package. We determined the significantly differentially expressed genes (p < 0.05) based on normalized counts and using the Benjamini-Hochburg correction (Benjamini & Hochburg, 1995) for multiple comparisons. We only retained genes with a -1 < log_2_ fold change > 1 in order to filter genes at a conservative threshold for differential expression based on our sample size (Schurch et al., 2016). This filtering was not necessary for either of the edgeR analyses because log_2_ fold changes exceeded this threshold for the differentially expressed genes and transcripts (-1.3 < log_2_ fold change > 1.4, in all cases).

We also compared the log_2_ fold change values (of treatment differences by mapped count) for each gene from the edgeR and DESeq2 gene-level analyses in a linear regression. This statistical test was performed in order to evaluate the degree of concordance between the two DGE analyses. Furthermore, we constructed a list of genes identified as differentially expressed by all three analyses, which were further evaluated for function as well as chromosomal location. These genes were also explored in STRING version 10.0 (string-db.org) to determine their protein-protein interactions (Snel et al., 2000; Szklarczyk et al., 2015).

Lastly, we performed an *a priori* test for DGE in edgeR on a small subset of nine genes encoding hormones and hormone receptors known to be involved in various aspects of reproductive functionality in male rodents. These genes are: steroidogenic acute regulatory protein (StAR), prolactin receptor (Prlr), luteinizing hormone/choriogonadotropin receptor (Lhgcr), inhibin (Inha), ghrelin (Ghrl), estrogen related receptor gamma (Essrg), estrogen related receptor alpha (Essra), androgen receptor (Ar), and activin receptor type-2A (Acvr2a). We retrieved the *Mus musculus* genomic sequences for these hormones and receptors from Ensembl (release 88: March 2017) and then executed BLASTn searches for the corresponding *Peromyscus eremicus* sequences in the testes transcriptome. The Ensembl gene identifiers (*Mus musculus*) corresponding to the *P. eremicus* transcripts were queried from the table of results produced by the edgeR DGE analysis to evaluate treatment differences in expression.

## Results

### Data and Code Availability

The testes transcriptome was assembled from a 45.8 million paired read data set. Additionally, there were 9-20 million paired reads for each of the 22 testes data sets used for the differential expression analysis **(Supplemental Table 1),** yielding 304,466,486 reads total for this analysis. The raw reads are available at the European Nucleotide Archive under study accession number PRJEB18655. All data files, including the testes un-annotated transcriptome, the dammit annotated transcriptome, and the data generated by the differential gene expression analysis (described below) are available on DropBox (https://www.dropbox.com/sh/ffr9xrmjxj9md1m/AACpxjQNn-Jlf25qNdslfRSCa?dl=0). These files will be posted to Dryad upon manuscript acceptance. All code for these analyses is posted on GitHub (https://github.com/macmanes-lab/testesDGE).

### Assembly of Testes Transcriptome

The performance of multiple transcriptome assemblies was evaluated thoroughly, and the selected optimized testes assembly met high quality and completeness standards, and it also contains relatively few contigs and has high read mapping rates **(Table 1).** Therefore, this transcriptome was used for our differential expression analyses. The transcriptome was also annotated, and the complete statistics for this dammit annotation are provided in **Table 1.**

**Table 1:**
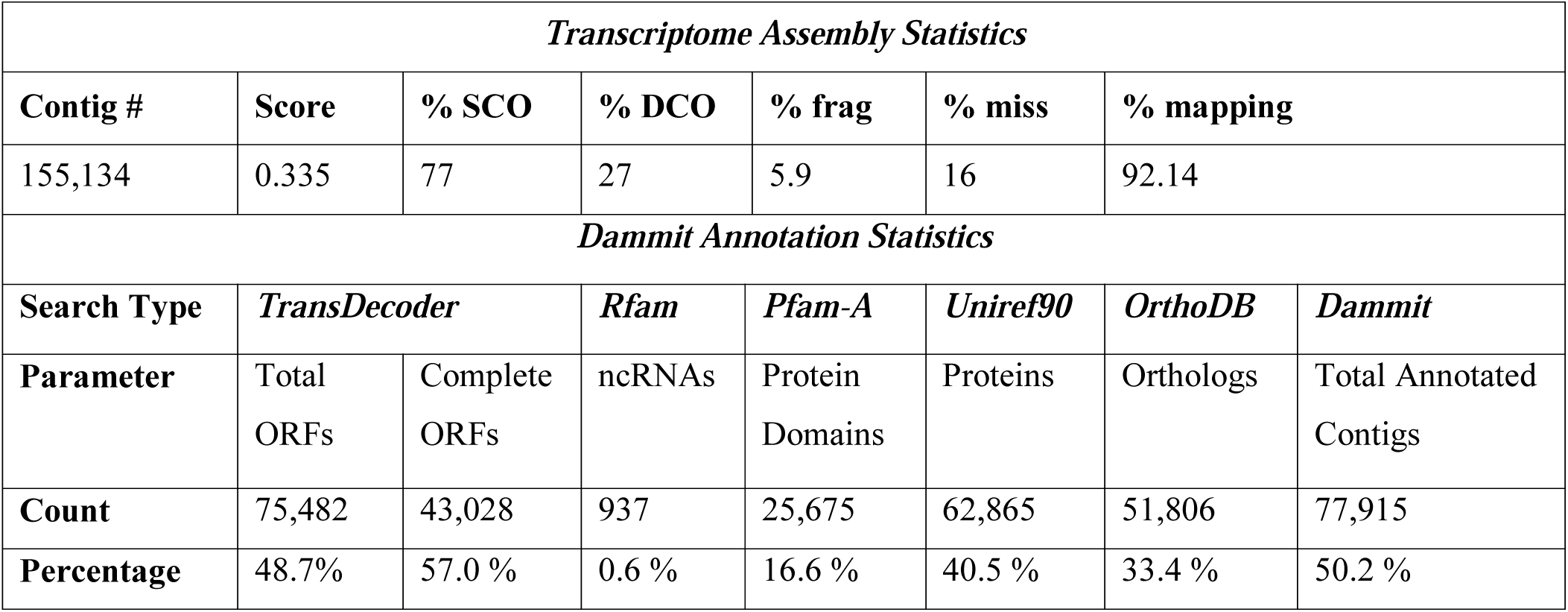
Transcriptome assembly (BinPacker CD-hit-est Transrate Corrected) performance metrics for: contig number, TransRate score (Score), BUSCO indices: % single copy orthologs (% SCO), % duplicated copy orthologs (% DCO), % fragmented (% frag), and % missing (% miss), as well as Salmon mapping rates (% mapping) for the optimized testes assembly. Dammit transcriptome assembly annotation statistics, including searches in the program TransDecoder for open reading frames (ORFs) and searches for homologous sequences in five databases: Rfam, Pfam-A, Uniref90, OrthoDB, and BUSCO. Percentages were calculated from the count number of each parameter divided by the total number of contigs in the transcriptome (155,134). The only exception to this calculation is for complete ORFs, which were calculated as a percentage of the total ORFs (75,482). The BUSCO results for the annotated assembly are not shown here as they are identical to those for the un-annotated assembly.

| Transcriptome Assembly Statistics |  |  |  |  |  |  |  |
| --- | --- | --- | --- | --- | --- | --- | --- |
| Contig # | Score | % SCO | % DCO | % frag | % miss | % mapping |  |
| 155,134 | 0.335 | 77 | 27 | 5.9 | 16 | 92.14 |  |
| Dammit Annotation Statistics |  |  |  |  |  |  |  |
| Search Type | TransDecoder |  | Rfam | Pfam-A | Uniref90 | OrthoDB | Dammit |
| Parameter | Total ORFs | Complete ORFs | ncRNAs | Protein Domains | Proteins | Orthologs | Total Annotated Contigs |
| Count | 75,482 | 43,028 | 937 | 25,675 | 62,865 | 51,806 | 77,915 |
| Percentage | 48.7% | 57.0 % | 0.6 % | 16.6 % | 40.5 % | 33.4 % | 50.2 % |

### Differential Gene and Transcript Expression Analyses

Salmon quasimapping rates of all read datasets to the assembly were sufficiently high (range: 81.46% - 87.02%; mean _WET_ =84.41; mean _DRY_ =83.81; **Supplemental Table 1),** indicating the successful generation of transcript count data for our differential expression analyses. The exact test performed for our gene-level analysis in edgeR indicated that fifteen genes reached statistical significance (after adjusting for multiple comparisons) for DGE between the WET and DRY treatment groups **(Supplemental Figure 1)**. Specifically, seven genes were more highly expressed in WET individuals, and eight genes were more highly expressed in DRY individuals **(Table 2).**

**Table 2:** EdgeR determined significantly differentially expressed genes by treatment group in *P*. *eremicus* testes. Of the 15 DGE, seven were significantly more highly expressed in WET mice (High in WET) and eight were more highly expressed in DRY mice (High in DRY).

| Ensembl ID | log<sub>2</sub>FC | logCPM | FDR | Gene ID | HIGH |
| --- | --- | --- | --- | --- | --- |
| ENSMUSG000000079019.2 | -4.354 | 1.650 | 5.82E-09 | Ins13 | WET |
| ENSMUSG000000054200.6 | -3.734 | 0.619 | 1.82E-06 | Ffar4 | WET |
| ENSMUSG000000026435.15 | -2.448 | 2.447 | 1.13E-03 | Slc45a3 | WET |
| ENSMUSG000000025020.11 | -2.231 | 1.770 | 1.13E-03 | Slit1 | WET |
| ENSMUSG000000031170.14 | -2.421 | 2.578 | 1.13E-03 | Slc38a5 | WET |
| ENSMUSG000000030830.18 | -2.180 | 1.666 | 3.37E-02 | Itgal | WET |
| ENSMUSG000000032554.15 | -2.066 | 3.287 | 4.85E-02 | Trf | WET |
| ENSMUSG000000001768.15 | 3.086 | 1.006 | 1.46E-07 | Rin2 | DRY |
| ENSMUSG000000025479.9 | 2.971 | 3.001 | 7.97E-05 | Cyp2e1 | DRY |
| ENSMUSG000000020427.11 | 2.681 | 3.887 | 1.13E-03 | Igfbp3 | DRY |
| ENSMUSG000000019997.11 | 2.314 | 3.235 | 1.13E-03 | Ctgf | DRY |
| ENSMUSG000000040170.13 | 1.951 | 0.753 | 1.72E-03 | Fmo2 | DRY |
| ENSMUSG000000023915.4 | 1.534 | 1.290 | 2.02E-02 | Tnfrsf21 | DRY |
| ENSMUSG000000052974.8 | 2.077 | 0.647 | 2.26E-02 | Cyp2f2 | DRY |
| ENSMUSG000000027901.12 | 2.492 | -0.620 | 4.78E-02 | Dennd2d | DRY |

We also performed an alternative transcript-level analysis using the referenced transcriptome mapped reads exclusively with edgeR. The exact test found 66 differentially expressed transcripts **(Supplemental Figure 2),** 45 of which were more highly expressed in the WET group, and 21 were more highly expressed in the DRY group **(Table 3).** 10 of these differentially expressed transcripts were consistent with differentially expressed genes from the edgeR DGE analysis. In addition, the significantly expressed transcripts without an Ensembl ID match (nine WET and nine DRY) were retrieved for performing an nt all species BLASTn search (http://blast.ncbi.nlm.nih.gov/Blast.cgi), and these results are in the supplementary materials.

**Table 3:**
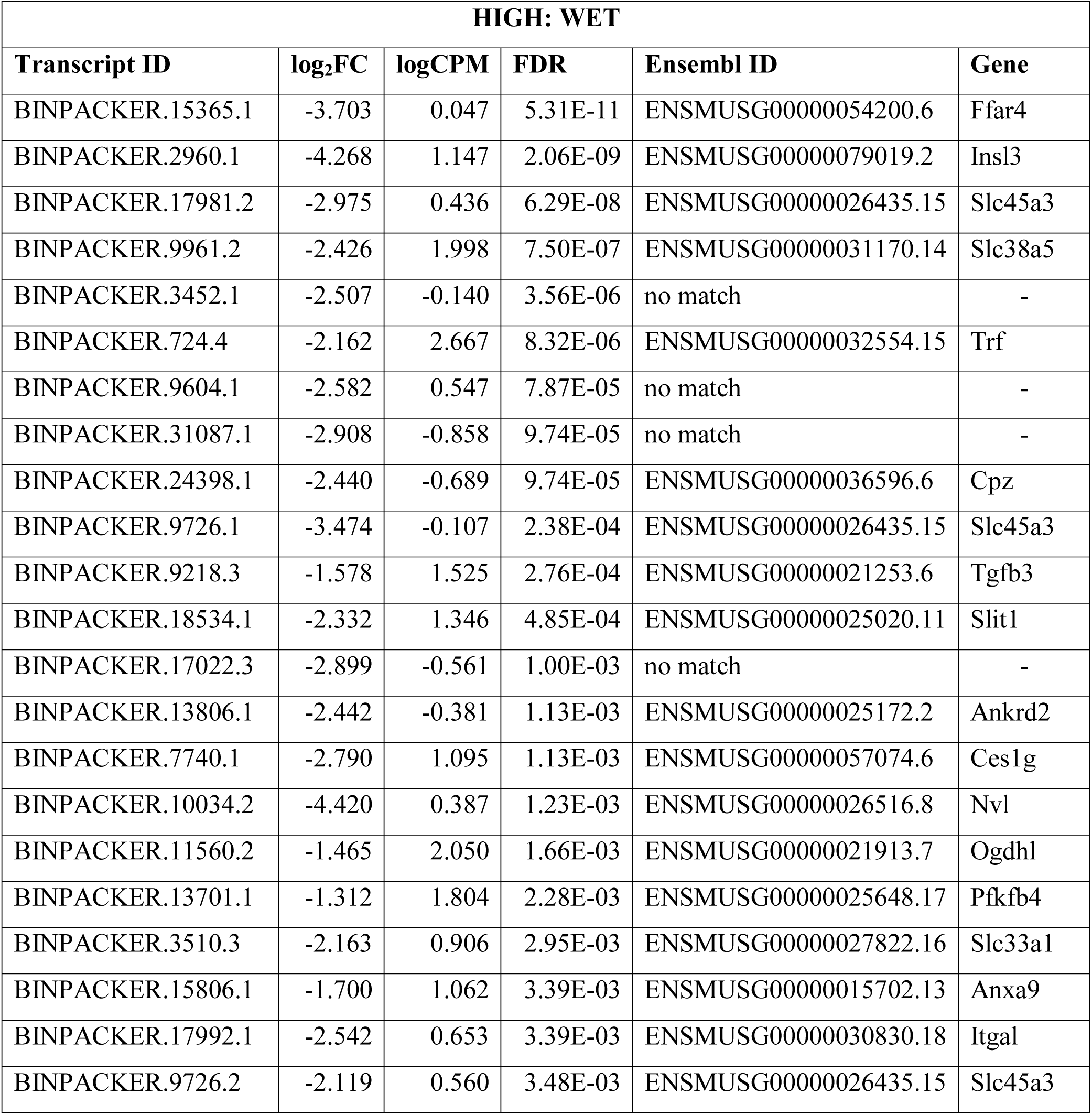
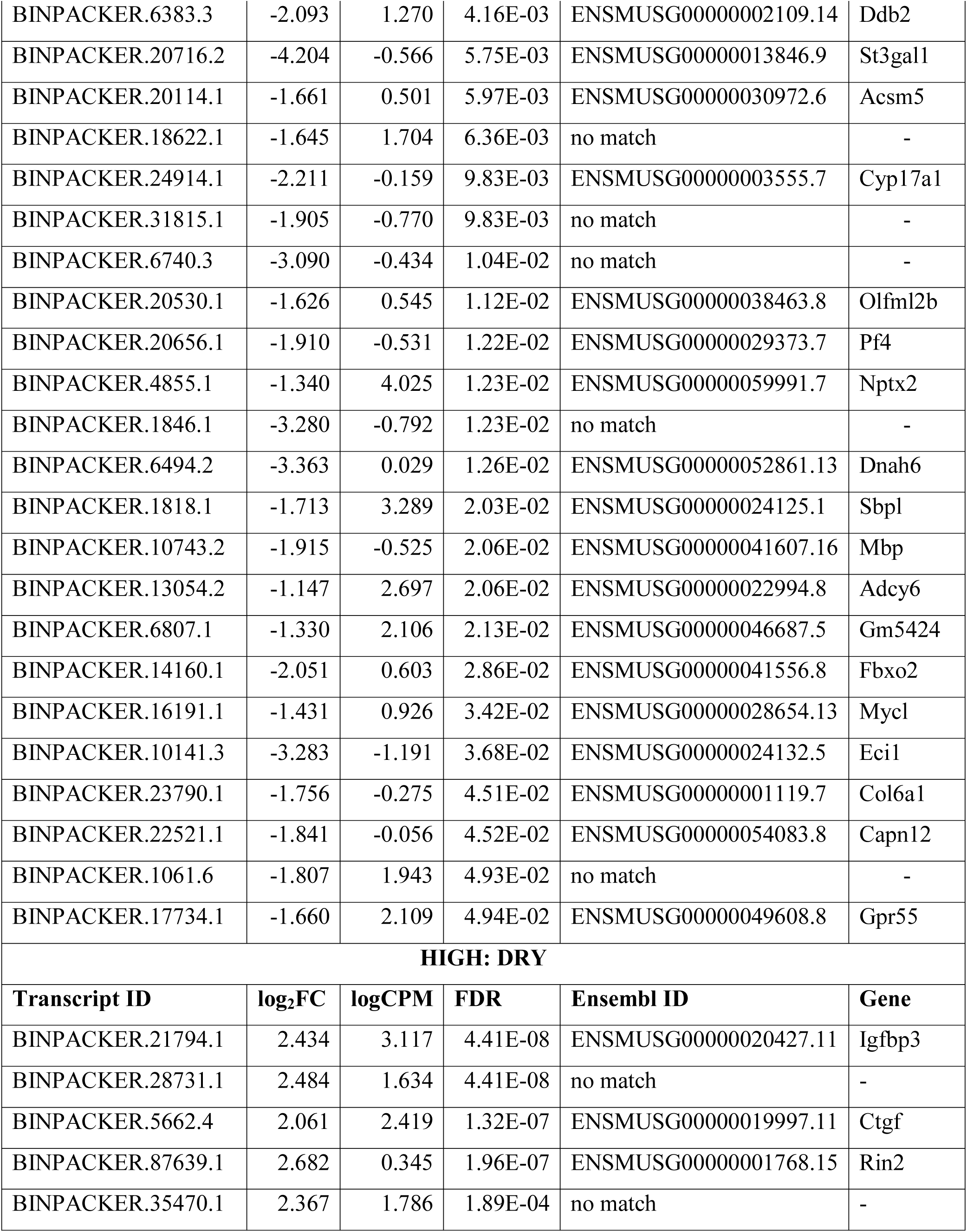
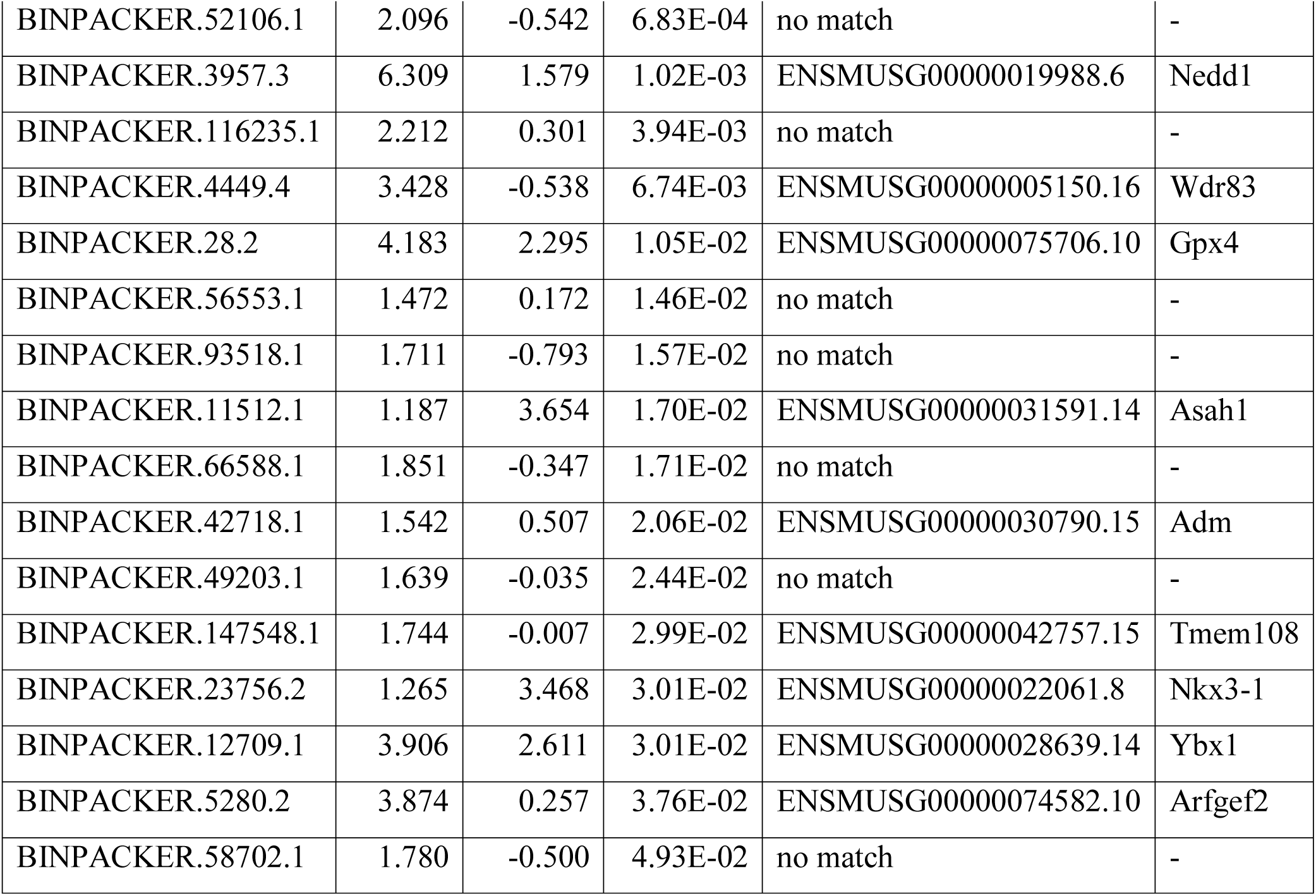
EdgeR determined significantly differentially expressed transcripts by treatment group in *P*. *eremicus* testes. Of the 66 total DTE, 45 were significantly more highly expressed in WET mice (High in WET) and 21 were more highly expressed in DRY mice (High in DRY). BLASTn matches to Ensembl IDs and corresponding Gene IDs.

The gene-level analysis conducted in DESeq2 yielded 215 significantly differentially expressed genes **(Supplemental Figure 3),** 67 of which were more highly expressed in the WET group, while 148 were highly expressed in the DRY group. However, only 20 of these genes remained when we filtered them with a -1 < log2 fold change > 1 to retain genes with a conservative threshold difference between treatment groups. This list of 20 genes yielded 16 genes more highly expressed in WET mice and four genes highly expressed in DRY mice **(Table 4).** Nine of these genes overlapped with those found to be significant in the previous two edgeR analyses.

**Table 4:** DESeq2 determined significantly differentially expressed genes by treatment group in *P*. *eremicus* testes. Of the 20 DGE with a -1 < log_2_ fold change > 1, 16 were significantly more highly expressed in WET mice (High in WET) and four were more highly expressed in DRY mice (High in DRY).

| Ensembl ID | baseMean | log<sub>2</sub>FC | p-adjusted | Gene ID | HIGH |
| --- | --- | --- | --- | --- | --- |
| ENSMUSG000000054200.6 | 8.77721485 | -2.2659204 | 1.24E-27 | Ffar4 | WET |
| ENSMUSG000000026435.15 | 38.7630267 | -2.2184407 | 1.16E-42 | Slc45a3 | WET |
| ENSMUSG000000079019.2 | 24.7158409 | -1.6454793 | 4.55E-13 | Insl3 | WET |
| ENSMUSG000000031170.14 | 42.2322119 | -1.6434261 | 6.64E-15 | Slc38a5 | WET |
| ENSMUSG000000038463.8 | 16.2605998 | -1.4619721 | 3.55E-12 | Olfml2b | WET |
| ENSMUSG000000030830.18 | 22.0478661 | -1.4358002 | 3.41E-10 | Itgal | WET |
| ENSMUSG000000032554.15 | 67.5197473 | -1.3762549 | 7.26E-10 | Trf | WET |
| ENSMUSG000000021253.6 | 31.2493344 | -1.3551661 | 7.02E-14 | Tgfb3 | WET |
| ENSMUSG000000030972.6 | 13.8934534 | -1.1709964 | 2.37E-07 | Acsn5 | WET |
| ENSMUSG000000059991.7 | 173.025492 | -1.1528314 | 5.12E-11 | Nptx2 | WET |
| ENSMUSG000000046687.5 | 44.9527785 | -1.0989949 | 8.31E-09 | Gm5424 | WET |
| ENSMUSG000000024125.1 | 101.5876 | -1.0962074 | 9.77E-06 | Sbpl | WET |
| ENSMUSG000000021913.7 | 46.5401886 | -1.0876018 | 8.70E-07 | Ogdhl | WET |
| ENSMUSG000000015702.13 | 27.7002506 | -1.0603879 | 1.95E-05 | Anxa9 | WET |
| ENSMUSG000000036596.6 | 6.6698922 | -1.0243046 | 9.04E-05 | Cpz | WET |
| ENSMUSG000000025172.2 | 13.2622565 | -1.0138171 | 0.00013318 | Ankrd2 | WET |
| ENSMUSG000000042757.15 | 14.5676529 | 1.00643936 | 0.00019556 | Tmem108 | DRY |
| ENSMUSG000000019997.11 | 64.49614 | 1.03331405 | 7.67E-05 | Ctgf | DRY |
| ENSMUSG000000020427.11 | 92.3763518 | 1.56656207 | 4.55E-13 | Igfbp3 | DRY |
| ENSMUSG000000001768.15 | 12.3794312 | 1.72433255 | 8.16E-16 | Rin2 | DRY |

To evaluate the correlation of log_2_ fold change results for each gene (Ensembl ID) from the two DGE analyses (EdgeR and DESeq2), we performed a regression of these log values, and they were significantly correlated **(Figure 1:** Adj-R^2^ = 0.6596; F(1,14214) = 2.754x10^4^; p < 2.2x10^−16^). This further demonstrates the concordance of the DGE analyses in these two software packages.

**Figure 1:**
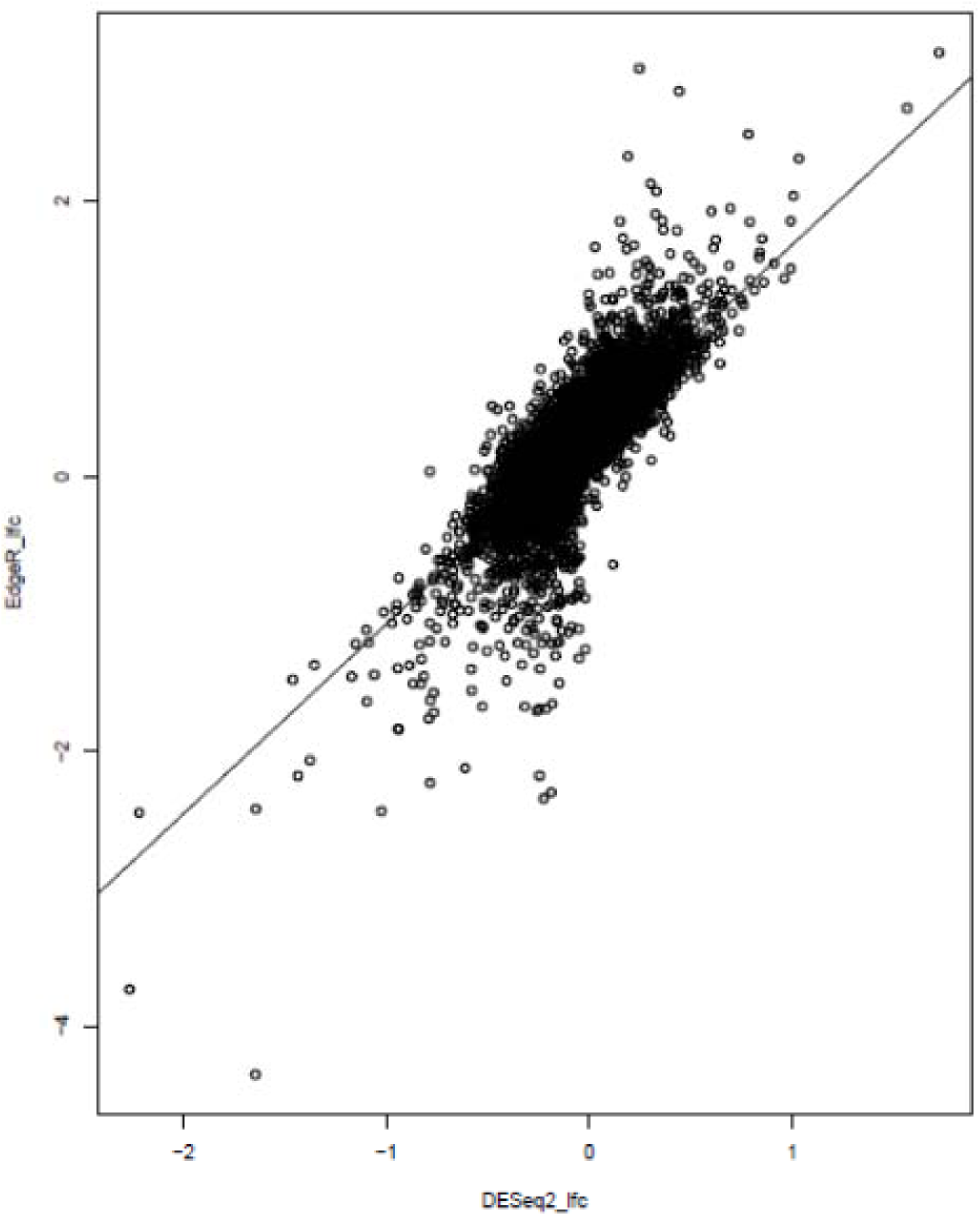
Correlation of log_2_ fold change results for all Ensembl ID gene matches from DESeq2 and edgeR DGE analyses (Adj-R^2^ = 0.6596; F(1,14214) = 2.754x10^4^; p < 2.2x10^−16^).

To evaluate the degree to which the three analyses produced concordant results, we generated a list of genes which were found to be significantly differently expressed by treatment across all three analyses **(Supplemental Table 2).** There were six genes that were consistently highly-expressed in the WET group and three genes that were highly-expressed in the DRY group. The six highly-expressed WET genes are Insulin-like 3 (Insl3), Free-fatty acid receptor 4 (Ffar4), Solute carrier family 45 member 3 (Slc45a3), Solute carrier family 38 member 5 (Slc38a5), Integrin alpha L (Itgal), and Transferrin (Trf). The three highly-expressed DRY genes are Ras and Rab Interactor 2 (Rin2), Insulin-like growth factory binding protein 3 (Igfbp3), and Connective tissue growth factor (Ctgf). Because the patterns of expression of these nine genes were corroborated by multiple methodologies, we are confident that they are differentially expressed between our treatments. Estimates of expression for these genes generated using the gene-level edgeR analysis are plotted in **Figure 2.**

**Figure 2:**
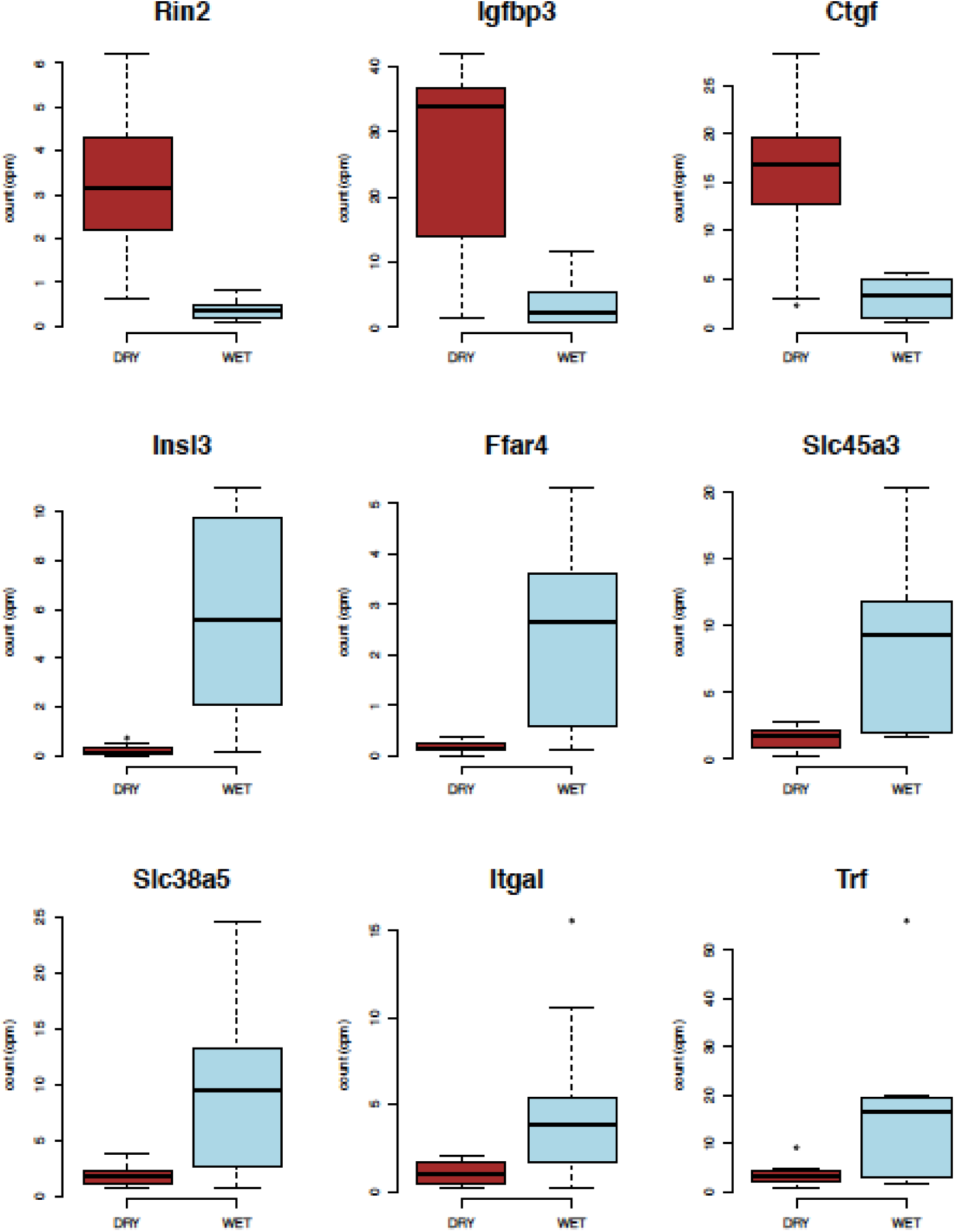
Box plots of edgeR analyzed differences in gene expression by treatment for the nine genes significantly differentially expressed in all three analyses. Counts per million (cpms) for both treatments (WET and DRY) are indicated.

The significantly differently expressed genes were evaluated for gene function and chromosomal location **(Table 5).** These genes occur throughout the genome; namely, they are located on different chromosomes. The diverse functions of each gene will be described below. In addition, we generated STRING diagrams (string-db.org) to view the protein-protein interactions for each of these nine genes (Snel et al., 2000; Szklarczyk et al., 2015).

**Table 5:** Functional information and chromosome (CHR) locations (*Mus musculus*) for the nine genes differentially expressed across all three analyses in *P. eremicus* testes by treatment group

| Gene Name | Gene ID | Gene Function | CHR | HIGH |
| --- | --- | --- | --- | --- |
| Insulin-like 3 | Ins13 | testicular function and testicular development | 8 | WET |
| Free-fatty acid receptor 4 | Ffar4 | metabolism and inflammation | 19 | WET |
| Solute carrier family 45 member 3 | Slc45a3 | sugar transport | 1 | WET |
| Solute carrier family 48 member 5 | Slc38a5 | sodium-dependent amino acid transport | X | WET |
| Integrin alpha L | Itgal | lymphocyte-mediated immune responses | 7 | WET |
| Transferrin | Trf | iron transport and delivery to erythrocytes | 9 | WET |
| Ras and Rab Interactor 2 | Rin2 | endocytosis and membrane trafficking | 2 | DRY |
| Insulin-like growth factor binding protein 3 | Igfbp3 | modulates effects of insulin growth factors | 11 | DRY |
| Connective tissue growth factor | Ctgf | fibrosis and extracellular matrix formation | 10 | DRY |

Slc38a5 and Slc45a3 are among the highly expressed genes in the WET group (they have lower expression in the DRY group); these two solute carriers are members of a large protein family that is responsible for cross-membrane solute transport (reviewed in Hediger et al., 2004; Hediger et al., 2013; Cesar-Razquin et al., 2015). Slc38a5 is involved sodium-dependent amino-acid transport, while Slc45a3 is purported to transport sugars (Vitavska and Wieczorek, 2013; Schiöth, et al., 2013; http://slc.bioparadigms.org/), thereby playing an important potential role in maintaining water balance via management of oncotic pressures. Slc38a5 **(Figure 3a)** has interactions with multiple additional solute carriers, including Slc1A5, Slc36A2, Slc36A3, and Slc36A4. Slc38a5 also has an interaction with disintegrin and metalloproteinase domain-containing 7 (Adam7), which is involved in sperm maturation and the acrosome reaction (Oh et al., 2005). In contrast, Slc45a3 **(Figure 3b)** does not have known protein interactions with other solute carriers; however, this protein does interact with steroidogenic acute regulatory protein (StAR), which is critical in steroidogenesis (Christenson and Strauss III, 2001). Notably, our *a priori* DGE analysis did not demonstrate treatment differences in expression for StAR.

**Figure 3:**
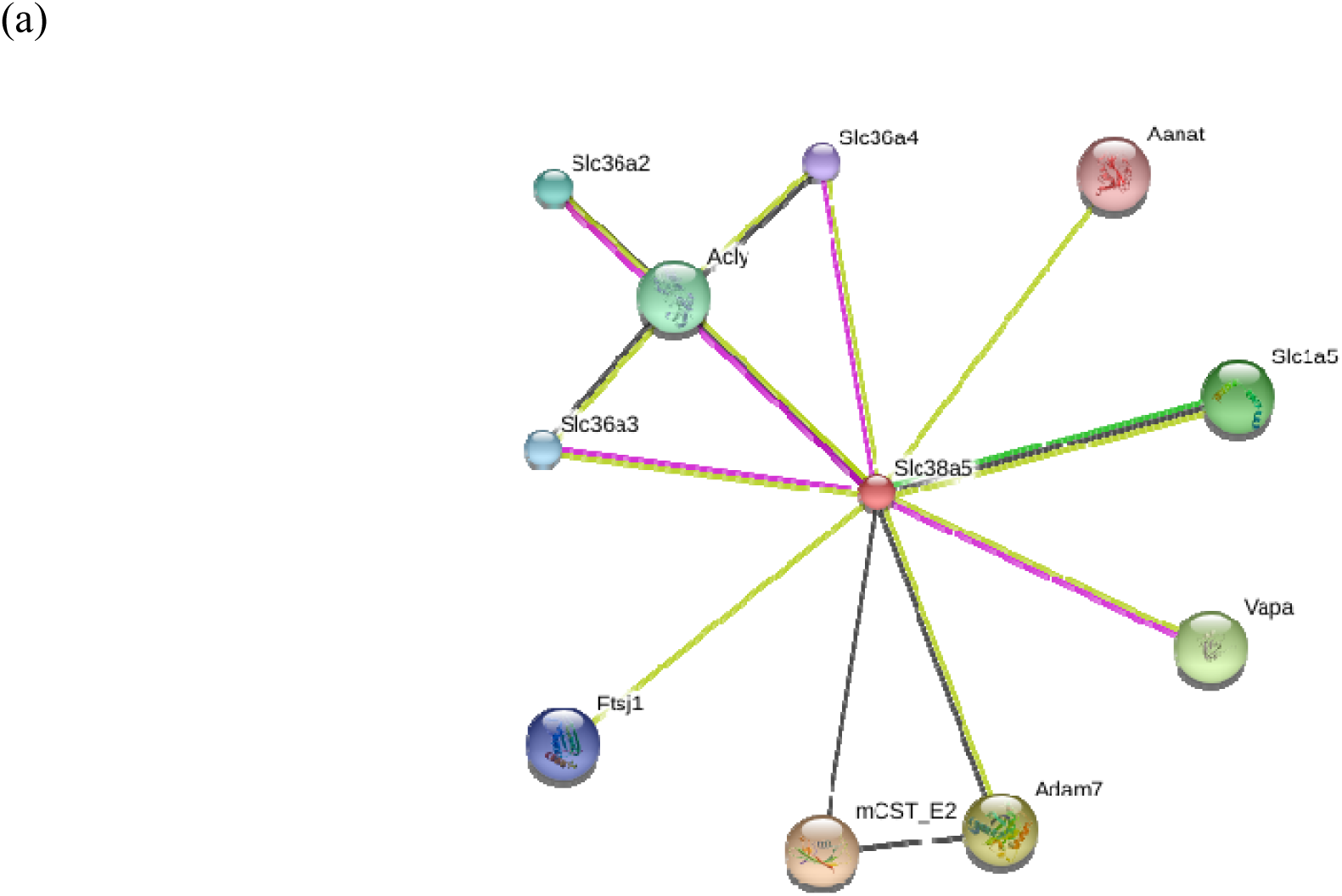
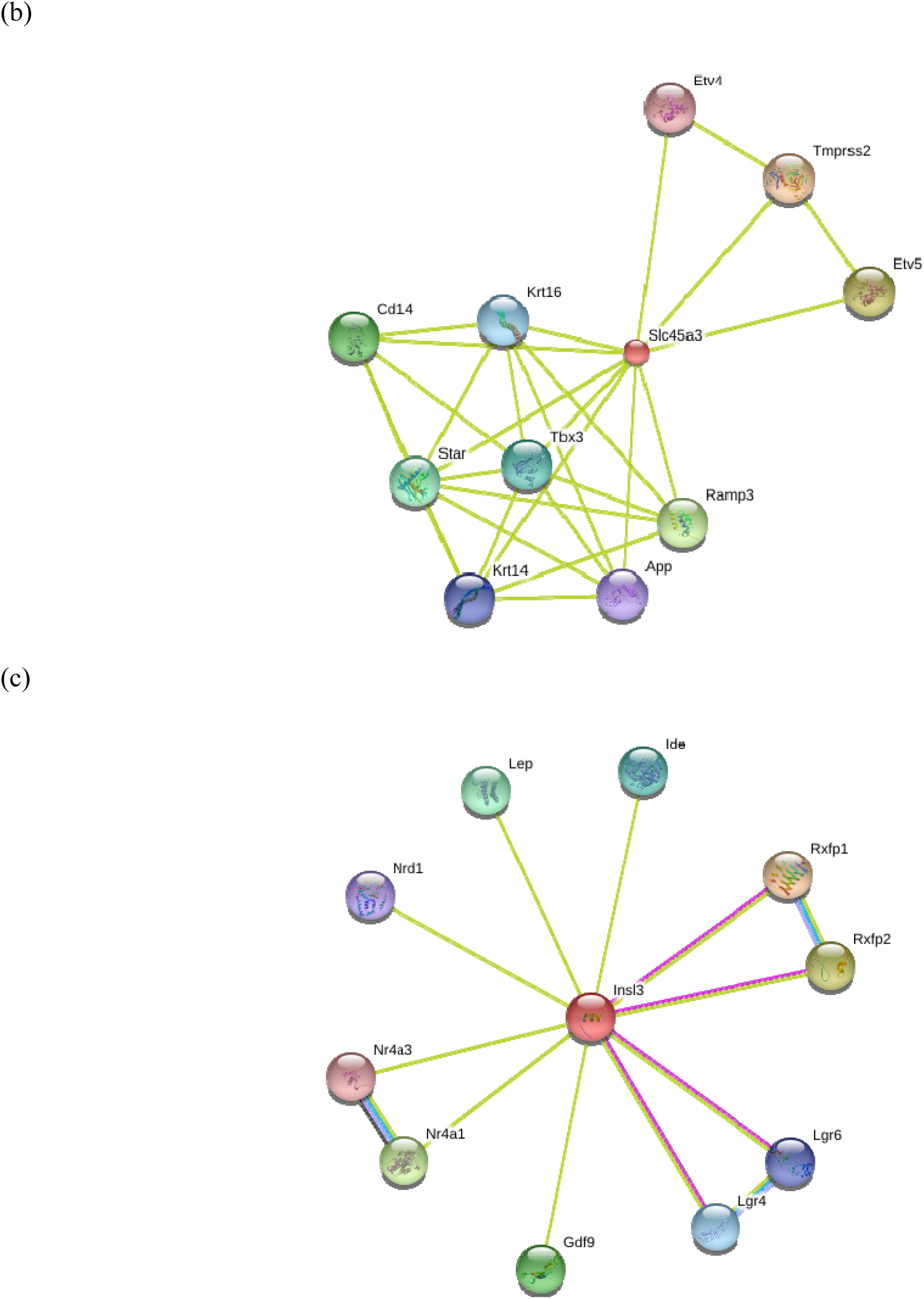
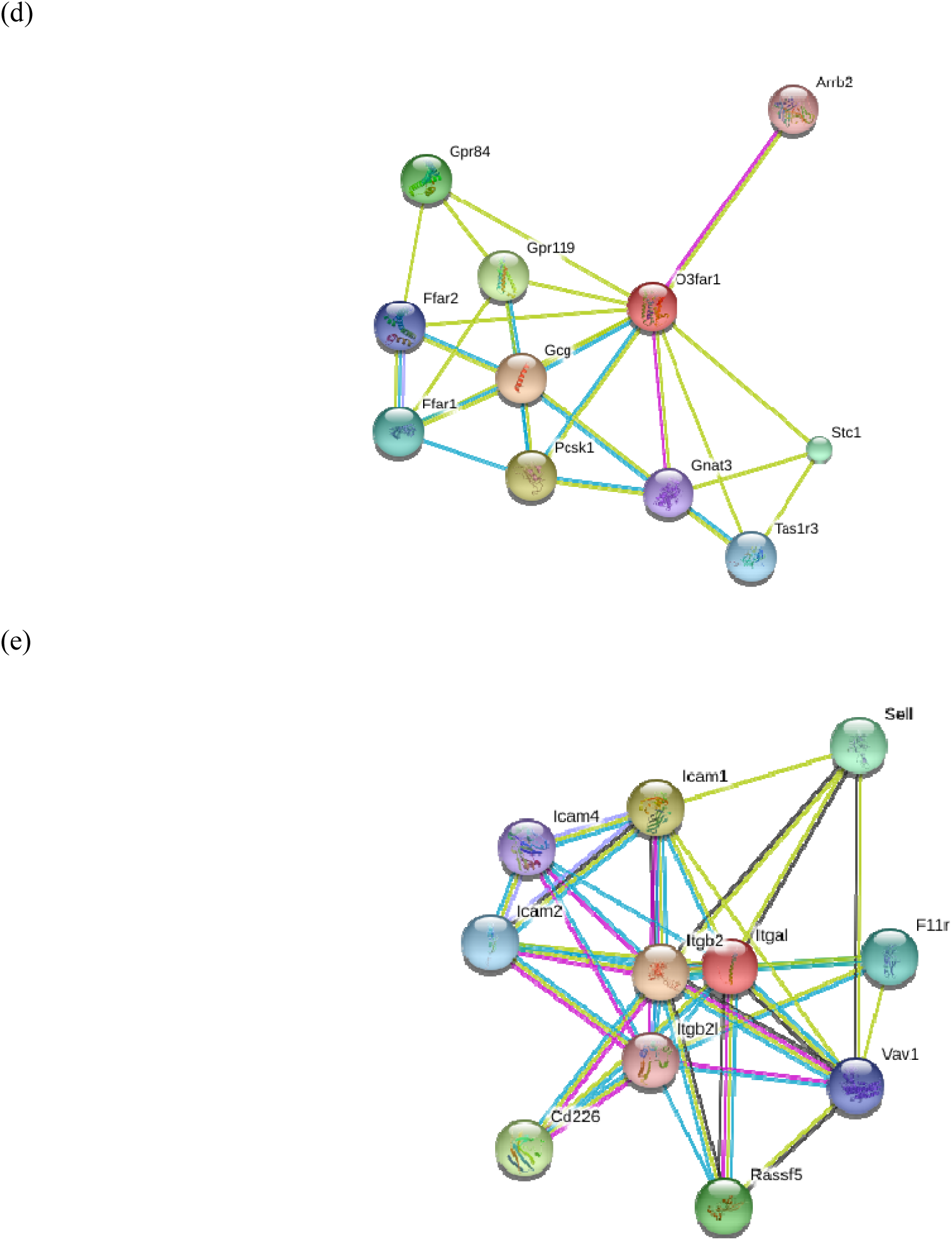
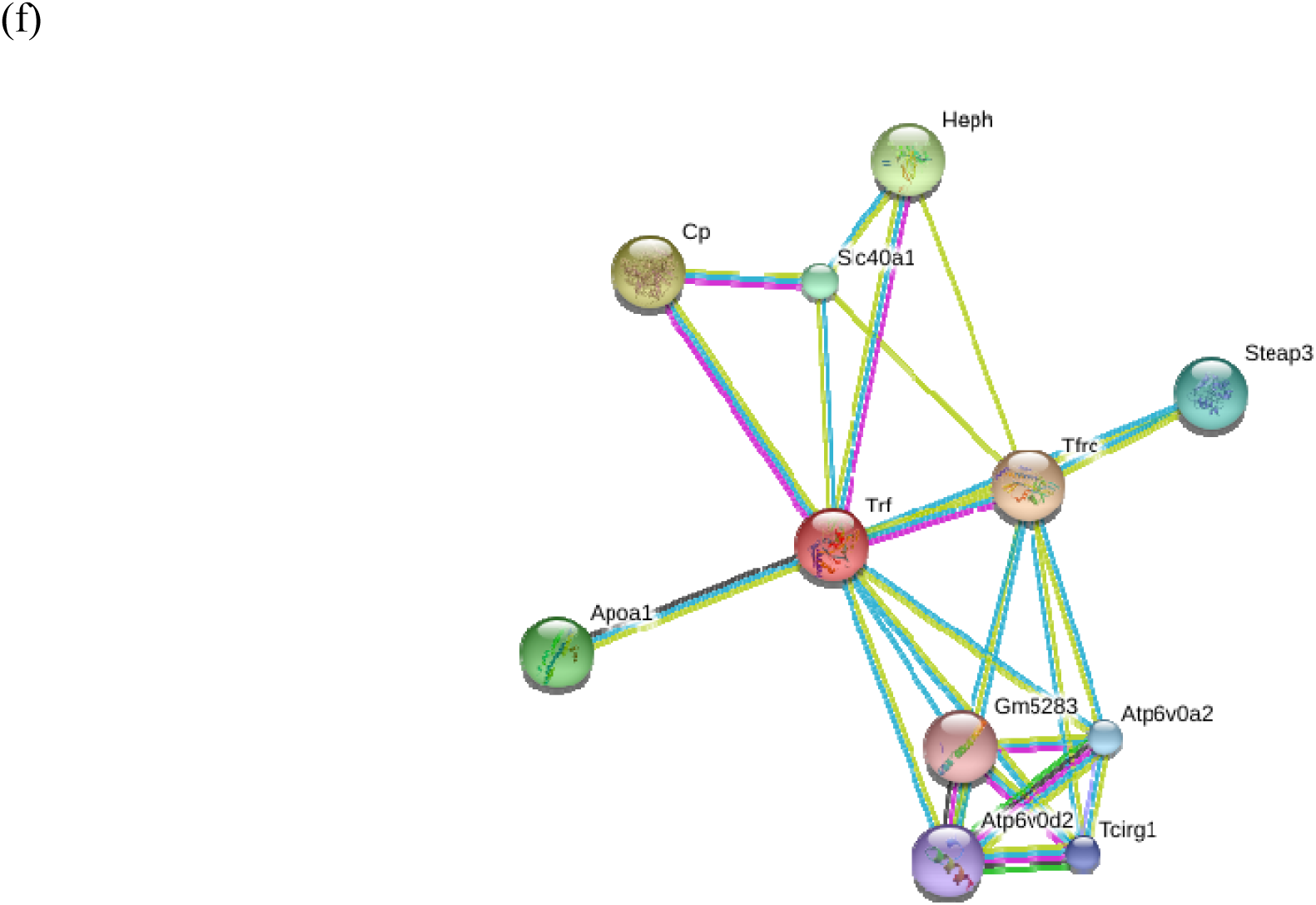
STRING diagrams of protein-protein interactions for genes significantly differentially expressed (highly expressed) in the WET treatment group. These six genes are (a) Slc38a5, (b) Slc45a3, (c) Insl3, (d) Ffar4 (also known as O3far1), (e) Itgal, and (f) Trf. Different colored circles stipulate different proteins interacting with the target proteins, small circles are proteins with unknown 3D structure, while larger circles are proteins with some degree of known or predicted 3D structure. Different colors of connecting lines represent different types of interactions between proteins. For fully interactive diagrams of the genes, view the provided links to string-db in the GitHub repository (StringDBlinks.md)

Insl3 was lower expressed in the DRY group, and this hormone purportedly regulates fertility in male and female mammals by preventing apoptosis of germ cells in reproductive organs of both sexes (Kawamura et al., 2004: Bathgate et al., 2012; Bathgate et al., 2013). In male rodents, Insl3 is critical to development by facilitating testicular descent, and it is also present in testes of adults, where it binds to relaxin family peptide receptor 2 (Rxfp2), also known as Lrg8 (Bathgate et al., 2012; Bathgate et al., 2013). Protein interaction data for Insl3 **(Figure 3c)** indicate that this hormone interacts with Rxfp2 and Rxfp1, as well as other proteins, including leptin (Lep), a pleiotropic hormone involved in reproduction, immunity, and metabolism (reviewed in Friedman, 2014).

Ffar4 was also down-regulated in the DRY group. Omega-3 fatty acid receptor 1 (O3Far1) is an alias of Ffar4, and it has roles in metabolism and inflammation (Moniri, 2016). This protein interacts with multiple other free fatty acid receptors and G-protein coupled receptors as well as Stanniocalcin 1 (Stc1) **(Figure 3d).** Stc1 is involved in phosphate and calcium transportation (Wagner and Dimattia, 2006); however, this protein’s functional role in mice remains enigmatic (Chang et al, 2005).

Another of the lower expressed DRY group genes is Itgal (also known as CDa11a), which has multifaceted roles in lymphocyte-mediated immune responses (Bose et al., 2014). Concordantly, the protein interactions with Itgal **(Figure 3e)** include numerous proteins integral to immunity, such as Intracellular adhesion molecules (specifically, ICAM1,2,4), which are expressed on the cell surface of immune cells and endothelial cells. Itgal is a receptor for these ICAM glycoproteins, which bind during immune system responses (reviewed in Albelda, Smith and Ward, 1994). However, an additional role of intercellular adhesion molecules has been proposed in spermatogenesis, whereby ICAMs may be integral to transporting non-mobile developing sperm cells through the seminiferous epithelium (Xiao, Mruk and Cheng, 2013).

The final gene with lower expression levels in the DRY treatment is Trf, which modulates the amount of free-iron in circulation and binds to transferrin receptors on the surface of erythrocyte precursors to deliver iron (reviewed in Gkouvastos Papanikolaou and Pantopoulos, 2012). Trf interacts with multiple proteins **(Figure 3f)** involved in iron transport and uptake, including Steap family member 3 (Steap3), hephaestin (Heph), cerulopslamin (Cp), Solute carrier protein 40 member 1 (Slc40A1), and several H+ ATPases. Furthermore, Trf is linked to apolipoprotein A-1 (Apoa1), which interacts with immunoglobulin in a complex named sperm activating protein (Spap) to activate the motility of sperm when it inhabits the female genital tract (Akerlof et al., 1991; Leijonhufvud, Akerlof and Pousette, 1997).

One of the highly expressed genes in the DRY group is Rin2, which is involved in endocytosis (reviewed in Doherty and McMahon, 2009) and membrane trafficking through its actions as an effector protein for the GTPases in the Rab family within the Ras superfamily (reviewed in Stenmark and Olkkonen, 2001). Rin2 protein-protein interactions **(Figure 4a)** include Ras related protein Rab5b and Rab5b, which are involved in vesicle transport as well as vasopressin-regulated water reabsorption. This mechanism for water reabsorption via Aquaporin 2 (Aqp2) in the kidney has been thoroughly reviewed by Boone and Deen (2008) and Kwon and colleagues (2013).

**Figure 4:**
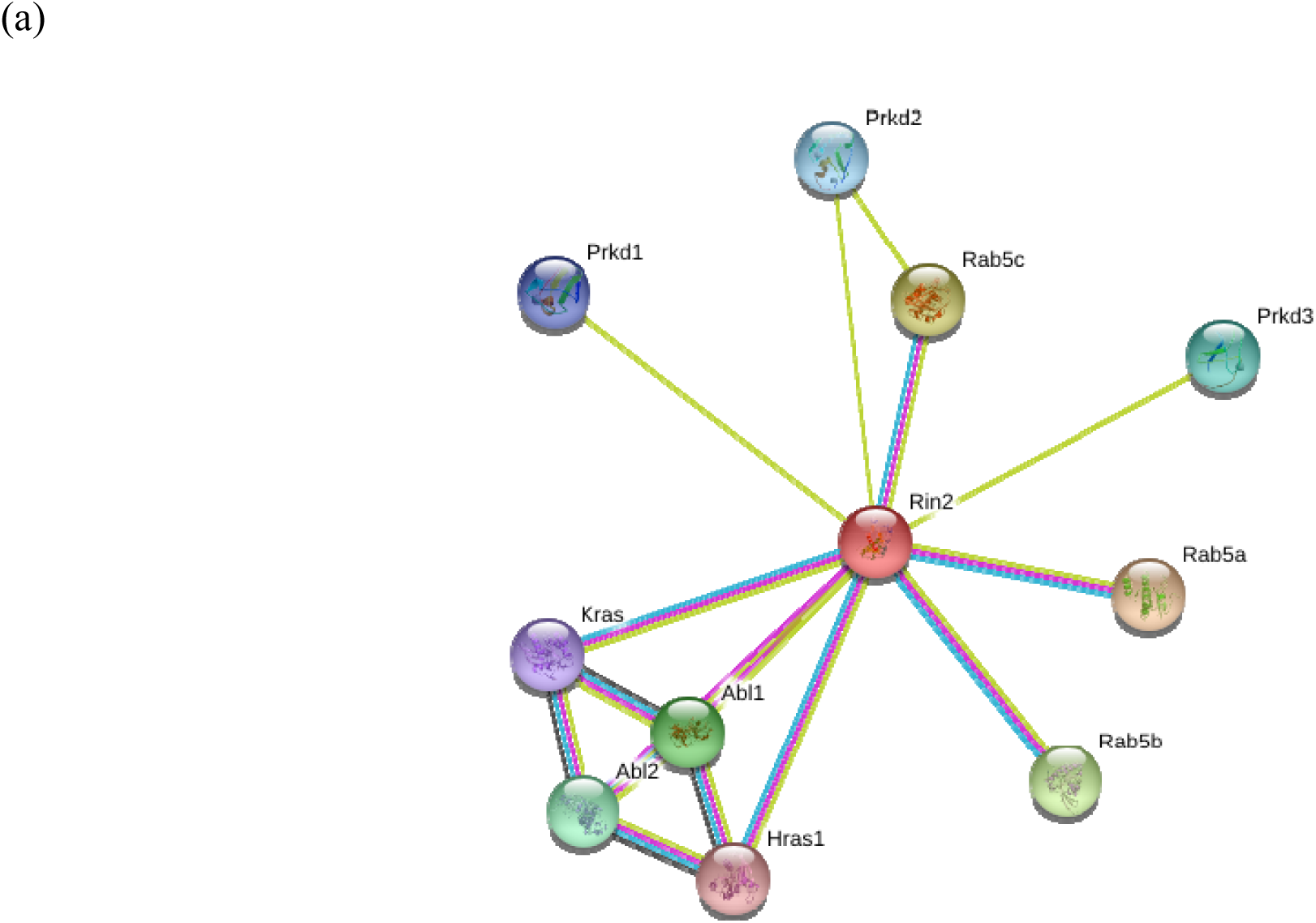
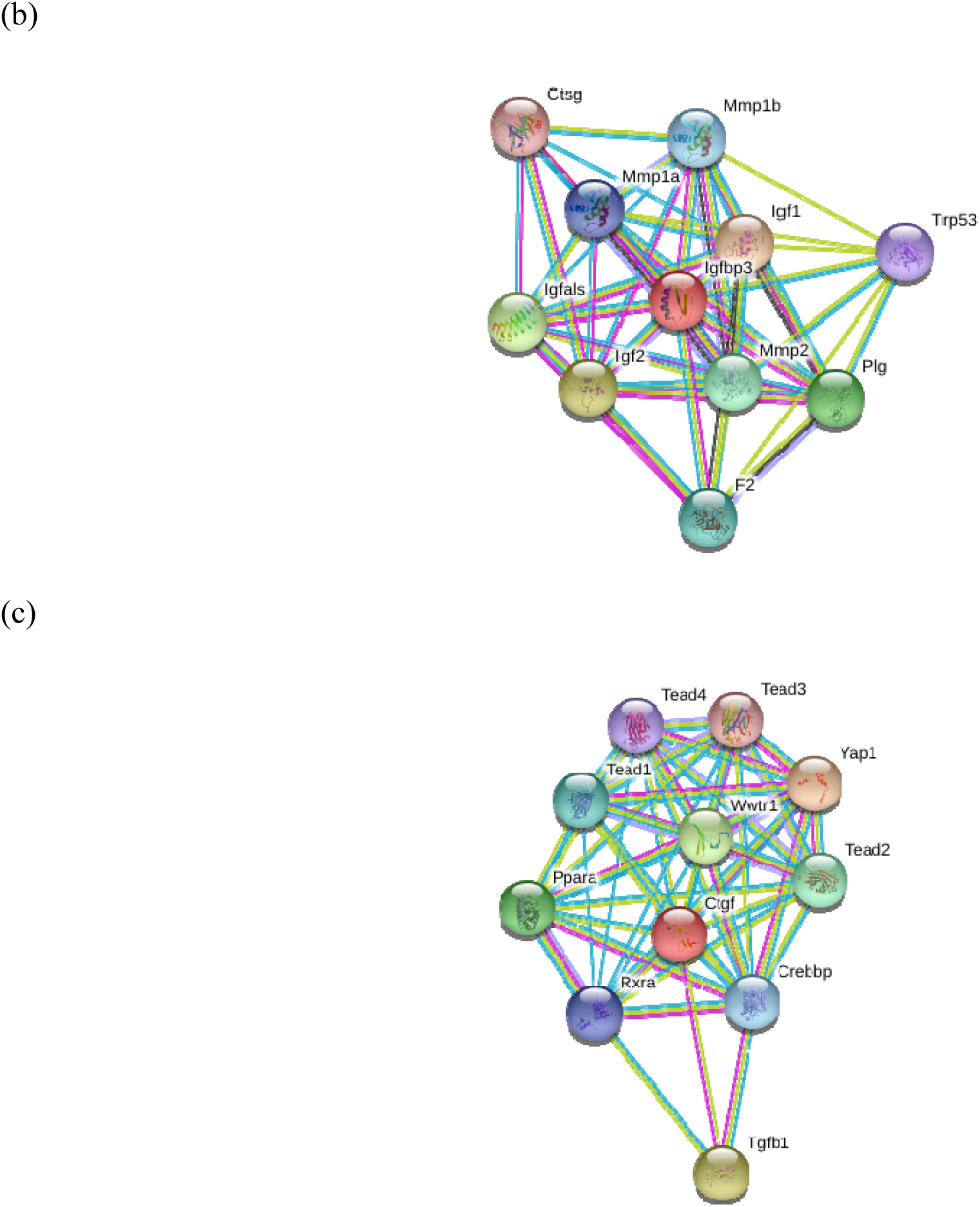
STRING diagrams of protein-protein interactions for genes significantly differentially expressed (highly expressed) in the DRY treatment group. These three genes are (a) Rin2, (b) Igfbp3, and (c) Ctgf. Different colored circles stipulate different proteins interacting with the target proteins small circles are proteins with unknown 3D structure, while larger circles are proteins with some degree of known or predicted 3D structure. Different colors of connecting lines represent different types of interactions between proteins. For fully interactive diagrams of the genes, view the provided links to string-db in the in the GitHub repository (StringDBlinks.md).

The second gene highly expressed in the DRY group is Igfbp3, which modulates the effects of insulin growth factors. Thus, the protein directly interacts **(Figure 4b)** with insulin growth factors 1 and 2 (Igf1, Igf2), which are responsible for increasing growth in most tissues (reviewed in le Roth 1997; Jones and Clemmons, 2008). Ctgf was also highly expressed in the DRY group, and this protein is responsible for increased fibrosis and extracellular matrix formation (Reviewed in Moussad and Brigstock, 2000). The protein interactions for Ctgf **(Figure 4c)** include many transcription activators in the Hippo signaling pathway, including multiple TEA domain transcription factors (Tead1, 2, 3 and 4), WW domain containing transcription regulator 1 (Wwtr1), as well as Yes-associated protein 1 (Yap1), which is responsible for both increasing apoptosis and preventing cell proliferation to mitigate tumor growth and control organ size (Reviewed in Pan, 2010).

The *a priori* edgeR DGE analysis for the genes encoding nine reproductive hormones and hormone receptors) did not reveal any statistically significant differences between the WET and DRY mice. The log fold change values and corresponding p-values for these genes are in the analysis posted on GitHub. The patterns for these genes by treatment are shown in **Figure 5.**

**Figure 5:**
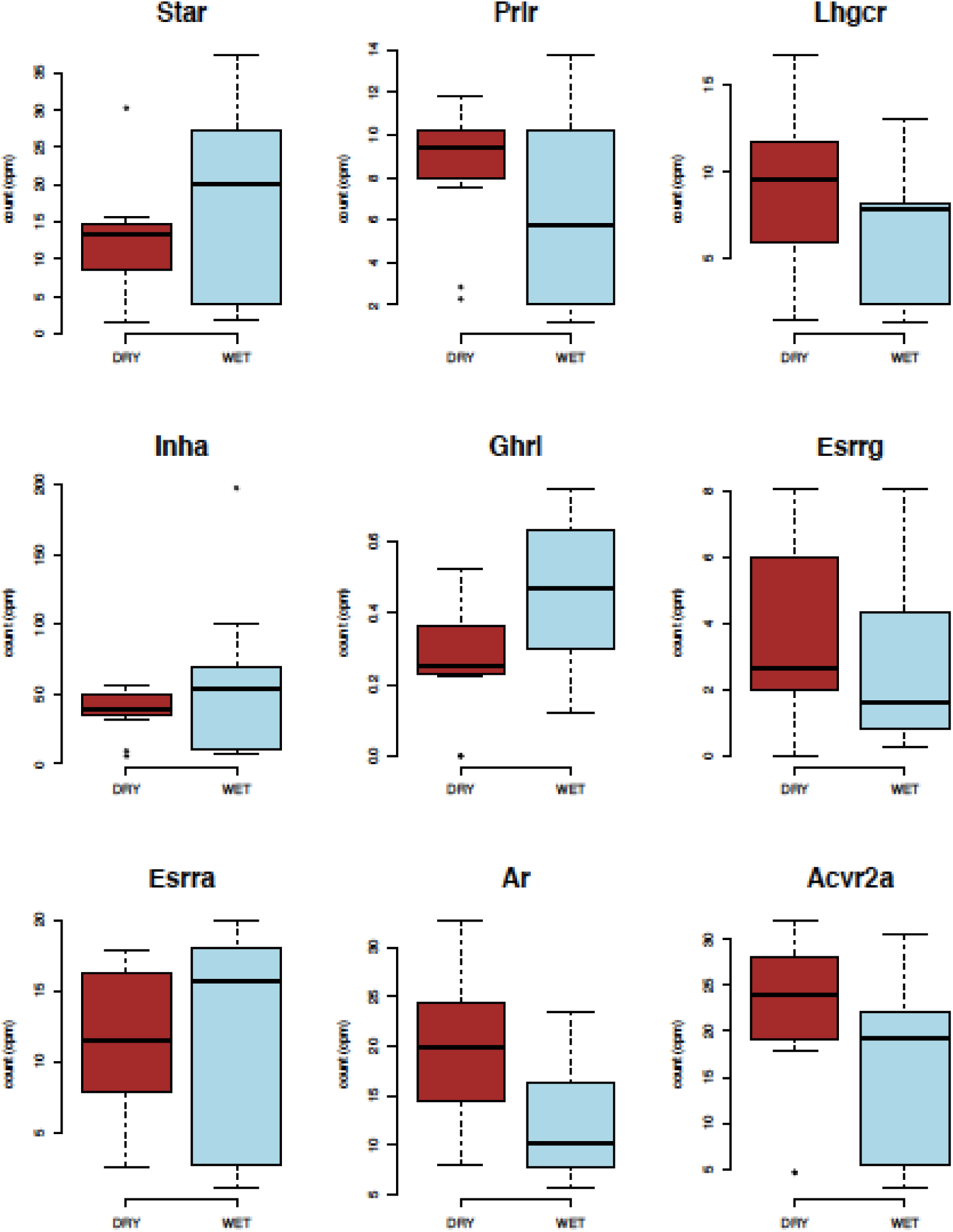
Box plots of edgeR analyzed differences in gene expression by treatment for the nine *a priori* tested reproductive hormone and hormone receptor genes. Counts per million (cpms) for both treatments (WET and DRY) are indicated.

## Discussion

This is the first study to evaluate gene expression levels of a reproductive tissue (testes) in response to acute dehydration in a desert-specialized rodent, *Peromyscus eremicus* (cactus mouse). Our results demonstrate differential expression of Insl3, which is a gene linked to reproduction, but not for a small subset of other reproductive hormone (and hormone receptor) genes. We also found expression differences in two solute carrier proteins, which is consistent with previous findings asserting the importance of this protein family for osmoregulation in desert rodents. Our findings lead us to hypothesize that reproductive function may be modified via Insl3 in acutely dehydrated mice. Any transcriptomic indication of potential reproductive modification in response to acute dehydration is surprising, given that this is not consistent with our understanding of *P. eremicus* as a desert specialist capable of breeding year-round in the wild. However, future studies must determine the physiological effects of decreased Insl3 expression on acutely dehydrated cactus mice. While acute dehydration is less common than chronic dehydration for desert mammals, given their ecology, it is a selective force they must overcome. Indeed, throughout much of the described range of the cactus mouse, rainfall events may occur several times per year. Cactus mice, and many other rodents, are known to rehydrate during these rainfall events (MacManes, *personal observation*). Following rehydration, cactus mice experience acute dehydration, followed by a steady state of chronic dehydration. The reproductive responses of cactus mice to these acute and chronic dehydration events are unknown; therefore, this study describes the transcriptomic effects of acute dehydration in testes.

Insl3, which is believed to be a hormonal regulator of fertility among mammals of both sexes, inhibits germ line apoptosis in the testes (Kawamura et al., 2004; Bathgate et al., 2012; Bathgate et al., 2013). Within adult rodent testes, luteinizing hormone (LH) stimulates expression of Insl3 in Leydig cells, and Insl3 binds to Lrg8 in seminiferous tubules, which results in inhibited apoptosis of germ-line cells, thus increasing their availability (Kawarmura et al., 2004). In addition, a study using murine Leydig cells demonstrated that Insl3 administration increased testosterone production (Pathirana et al., 2012). The precise mechanistic role of ISNL3 in modulating fertility is still being elucidated; however, researchers assert that this hormone is an important regulator of fertility in males and females (*reviewed in* Bathgate et al., 2012). Indeed, recent research has investigated the utility of Insl3 as an indicator of mammalian fertility (e.g. *in humans:* Kovac and Lipshultz, 2013; *in bulls*: Pitia et al., 2016). Insl3 is also critical for the first phase of testicular descent, the transabdominal phase, which occurs during fetal development in rodents; but Insl3 does not appear to be involved in the inguinoscrotal phase which happens in sexually immature or inactive male rodents (reviewed in Hutson et al., 2015). Lower Insl3 expression in the testes of acutely dehydrated mice leads us to suggest that fertility may be attenuated due to acute water deprivation. However, future work characterizing the functional consequences of Insl3 down-regulation, including direct measurements of sperm numbers and function, is needed to causatively demonstrate reproductive attenuation. Specifically, does the number or quality of sperm decrease, and does this decrease reduce the probability of successful fertilization? Moreover, what are the temporal dynamics of reproductive suppression? Logically, species with core reproductive functions that are suppressed by dehydration seem likely to be rapidly outcompeted by others lacking such limitations. Given this assertion, research characterizing the reproductive correlates of chronic dehydration is a logical extension of this work, although doing so is beyond the scope of this study.

Solute carrier proteins, specifically Slc45a3 and Slc38a5, are downregulated in acute dehydration. These genes are part of a large family essential for transferring solutes across membranes (reviewed in Hediger et al., 2004; Hediger et al., 2013; Cesar-Razquin et al., 2015). Another member of this family, Solute carrier family 2 member 9 (Slc2A9), has been found to be undergoing positive selection in studies on kidney transcriptomes of cactus mouse (MacManes & Eisen, 2014) and of other desert rodents (Marra, Romero & DeWoody, 2014). Our previous work with the male reproductive transcriptome of cactus mouse found evidence for positive selection in two additional solute carrier proteins: Slc15a3 and Slc47a1 (Kordonowy and MacManes, 2016). A recent differential gene expression study in cactus mouse kidneys found that Slc2A1 and Slc8A1 also showed responses to acute dehydration (MacManes, 2017). Therefore, our current findings that two solute carrier proteins are lower expressed in the DRY treatment group is consistent with previous research in the kidney and male reproductive transcriptomes for this species. This leads us to further support the hypothesis originally proposed by Marra, Romero & DeWoody (2014) that this protein family is intrinsic to osmoregulation in desert rodents. Indeed, the findings of MacManes and Eisen (2014), Kordonowy and MacManes (2016), and MacManes (2017) also lend support to the essential role of solute carrier proteins for maintaining homeostasis in the desert specialized cactus mouse.

In addition to their well characterized role in the maintenance of water and electrolyte balance, the differential expression of solute carrier proteins may have important reproductive consequences, particularly as they relate to hormone secretion. Indeed, the interaction between Slc38a5 and Adam7 is relevant, because Adam7 is involved in sperm maturation and the acrosome reaction (Oh et al., 2005). Furthermore, the protein-protein interactions between Slc45a3 with StAR and between Insl3 and Lep are of particular interest because both StAR and Lep are integral to reproduction, as well as to homeostasis (reviewed in Christenson and Strauss III, 2001; Anuka et al., 2013; Friedman, 2014; Allison and Myers, 2014). However, our a priori DGE analysis evaluating StAR, and other reproductive hormones, did not show evidence of expression changes. Thus, the protein interactions with reproductive implications are not restricted to solute carrier proteins. The protein relationships between Itgal and intercellular adhesion molecules are also noteworthy with respect to research hypothesizing an integral role for ICAMs in spermatogenesis (Xiao, Mruk and Cheng, 2013). Furthermore, Trf is linked to Apoa1, which is a critical component of sperm activating protein (Akerlof et al., 1991; Leijonhufvud, Akerlof and Pousette, 1997). While the relationship between these differentially expressed genes and the hormones involved in reproductive function are currently poorly-characterized, our findings that genes integral to sperm development and activation interact with genes differentially expressed in acute dehydration may indicate that, contrary to our expectations, acute dehydration is linked to reproductive modulation in the cactus mouse. However, functional studies will be necessary to elucidate the connection between these genes and physiological responses to dehydration. This is particularly important because many hormones have pleotropic effects, and further mechanisms of action unrelated to reproduction may be elucidated for these proteins in *Peromyscus eremicus*.

In contrast to genes that are down-regulated in dehydration, the genes that were upregulated in the DRY group are known to be responsible for water homeostasis and cellular growth. The significance of Rin2 is notable, because this protein is an effector for Rab5, which as a GTPase involved in vasopressin-regulated water reabsorption, a critical homeostatic process mediated through the Aqp2 water channel in kidneys (Boone and Deen, 2008; Kwon et al., 2013). It is not surprising that genes in addition to solute carrier proteins, which are implicated in alternative processes for water homeostasis, are differentially expressed in response to water limitation. The other two genes that are up-regulated in the DRY treatment are indicative of modulated growth due to water limitation. Specifically, Igfb3 interacts directly with insulin growth factors responsible for tissue growth (le Roth 1997; Jones and Clemmons, 2008), and Ctgf is linked with numerous transcription factors in the Hippo signaling pathway, which modulates apoptosis, proliferation and organ size control (Pan, 2010).

To complement our male centric research, future studies should evaluate dehydration induced gene expression differences in female reproductive tissues, particularly in the uterus and ovaries during various reproductive stages. Indeed, given that the physiological demands of reproduction are purportedly greater in females, though this is controversial, (Bateman’s Principle: *proposed in* Bateman, 1948; *addressed in* Trivers, 1972; *reviewed in* Knight, 2002; *tested in* Jones et al., 2002; 2005; Collet et al., 2014), we would expect to see a greater degree of reproductive suppression in females. While such work is beyond the scope of this manuscript, we hope that future research will evaluate female Cactus mouse reproductive responses to dehydration.

Our findings are pertinent to physiological research in other desert-rodents showing reproduction suppression in response to water limitation (*reviewed in* Bales and Hostetler, 2011), specifically, in male and female Mongolian gerbils (Yahr and Kessler, 1975) and female hoping mice (Breed, 1975). The integral role of water as a reproductive cue for desert-rodents has also been demonstrated in water-supplementation studies (*reviewed in* Bales and Hostetler, 2011; Christian, 1979) as well as research on the effects of desert rainfall (Breed and Leigh, 2011; Henry and Dubost, 2012; Sarli et al., 2015; Sarli et al., 2016). Thus, Schwimmer and Haim (2009) asserted that reproductive timing is the most evolutionarily important adaptation for desert rodents. Furthermore, desert rodent research supporting a dehydration driven reproductive suppressive pathway mediated by arginine vasopressin (*reviewed in* Schwimmer and Haim, 1999; tested in Tahri-Joutei and Pointis, 1988a; 1988b; Shanas and Haim, 2004; Wube et al., 2008; Bukovetzky et al., 2012a; Bukovetzky et al., 2012b) is somewhat analogous to our study linking decreased Insl3 expression in testes with dehydration, in that both findings represent non-traditional hormonal modulation of reproduction. We propose that future studies thoroughly explore physiological consequences for non-traditional hormonal pathways in response to dehydration in desert rodents, as well as well-established reproductive modulatory hormones in the hypothalamic-pituitary-gonadal axis.

Emerging from this work is a hypothesis related to the reproductive response to water stress in the cactus mouse, and perhaps other desert rodents. Specifically, we hypothesize that acute dehydration may be related to reproductive mitigation; however, we hypothesize that chronic dehydration is not. Indeed, it is virtually oxymoronic to suggest that chronic dehydration, which is the baseline condition in desert animals, has negative consequences for reproductive success. Indeed, desert rodents dynamically respond to water-availability to initiate and cease reproductive function. Generating an integrative, systems-level understanding of the reproductive responses to both acute and chronic dehydration across desert-adapted rodent is required for testing our hypothesis. While understanding the renal response to dehydration is critical for making predictions about survival, understanding the reproductive correlates is perhaps even more relevant to evolutionary fitness. This study, to the best of our knowledge, is the first to describe the reproductive correlates of water-limitation in the cactus mouse, and the first to use a differential gene expression approach to evaluate reproductive tissue responses to drought. Furthermore, this study contributes to a research aim to determine whether novel physiological reproductive adaptations are present in male Cactus mouse (Kordonowy and MacManes, 2016). Developing a comprehensive understanding of reproductive responses to drought, and also the mechanisms underlying potential physiological adaptations, is necessary if we are to understand how increasing environmental variability due to climate change may modify the distribution of extant organisms.

## Conclusions

The genetic mechanisms responsible for physiological adaptations for survival and reproduction in deserts remain enigmatic. Desert rodent research has focused primarily on physiological adaptations related to survival, specifically on renal adaptations to combat extreme water-limitation. In contrast, while previous studies have investigated reproductive effects of water-limitation in desert rodents, the underlying mechanisms for physiological adaptations for reproduction during acute and chronic dehydration are unknown. Furthermore, ours is the first study to evaluate reproductive transcriptomic responses to water limitation in a desert-rodent, the cactus mouse. To this end, we characterized the reproductive correlates of acute dehydration in this desert-specialized rodent using a highly replicated RNAseq experiment. In contrast to expectations, we describe a potential signal of reproductive modulation in dehydrated male cactus mouse testes. Specifically, dehydrated mice demonstrated significantly lower expression of Insl3, which is a canonical regulator of fertility (and testes descent). Lower expression was also found in Slc45a3 and Slc38a5, lending further credence to the important role of solute carrier proteins for osmoregulation in the cactus mouse. While the low number of differentially expressed genes between acutely dehydrated and control mice might otherwise have suggested that this species is relatively unaffected by acute water-limitation, the diminished expression of Insl3 in dehydrated mice leads us to propose that acute dehydration may compromise reproductive function via decreased fertility. Indeed, we hypothesize that non-traditional reproductive hormone pathways, such as those involving Insl3 or AVP (which has elicited suppressive reproductive responses in other desert rodent research), warrant further investigation in studies evaluating the reproductive effects of acute and chronic dehydration. Although future research must experimentally evaluate the potential functional relationship between Insl3 expression pattern and reproductive function and fertility, our findings that acute-dehydration alters Insl3 expression may be concerning, particularly with respect to global climate change. Climate change driven increased variabilities in weather patterns may result in a greater frequency of acute water-stress, which could result in reduced reproductive function for the cactus mouse. In addition, because global climate change is predicted to shift habitats toward extremes in temperature, salinity, and aridity, and to alter species ranges, an enhanced understanding of the reproductive consequences of these changes, and of the potential for organisms to rapidly adapt, may enable us to effectively conserve innumerable species facing dramatic habitat changes.

## List of abbreviations

(DGE): Differential Gene Expression
(DTE): Differential Transcript Expression
(DRY): Dehydrated Treatment Group
(WET): Control Treatment Group

## Declarations

### Ethics approval and consent to participate

All animal care procedures were conducted in accordance with University of New Hampshire Animal Care and Use Committee guidelines (protocol number 130902) and guidelines established by the American Society of Mammalogists (Sikes et al., 2016).

### Consent for publication

Not applicable.

### Availability of data and materials

The raw reads are available at the European Nucleotide Archive under study accession number PRJEB18655. All data files, including the testes un-annotated transcriptome, the dammit annotated transcriptome, and the data generated by the differential gene expression analysis (described below) are available on DropBox (https://www.dropbox.com/sh/ffr9xrmjxj9md1m/AACpxjQNn-Jlf25qNdslfRSCa?dl=0). These files will be posted to Dryad upon manuscript acceptance. All code for these analyses is posted on GitHub (https://github.com/macmanes-lab/testesDGE).

### Authors’ Contributions

LK and MDM both contributed to the data collection, data generation, bioinformatics, analyses, interpretation, and writing of this manuscript.

## Acknowledgments

We would like to acknowledge the MacManes laboratory, including graduate student Andrew Lang, and the undergraduate students in the laboratory. We also acknowledge Dr. Paul Tsang for advice on the reproductive endocrinology described in the discussion.

---

**Supplemental Table 1:**
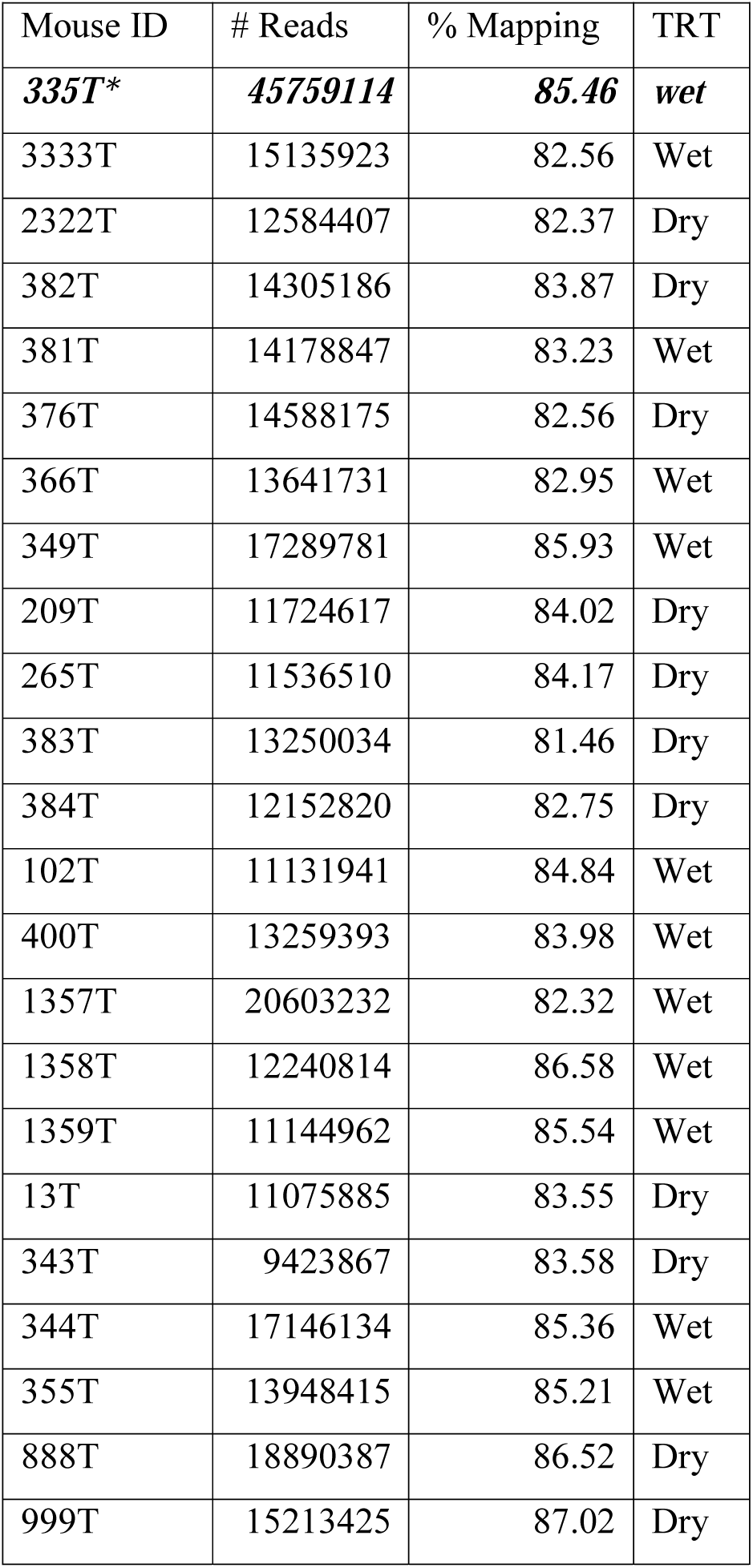
Testes read data statistics, including sample identification (Mouse ID), number of reads (# Reads), percent reads mapped to transcriptome (% Mapping), and treatment group (TRT). Mouse ID 335T* is the dataset which was used to assemble the testes transcriptome; therefore, these reads were not used for the differential expression analysis.

**Supplemental Table 2:** Significantly differentially expressed genes identified in the three analyses (DGE in edgeR, DTE in edgeR, and DGE in DESeq2) by treatment group in *P. eremicus* testes. Of the 34 different genes which were more highly expressed in WET mice, six were significant across all three analyses (Gene IDs are italicized). Of the 17 genes which were more highly expressed in DRY mice, three were significant across all three analyses (Gene IDs are italicized).

| **HIGH: WET** |  |  |  |
| --- | --- | --- | --- |
| **Gene ID** | **DGE edgeR** | **DTE edgeR** | **DGE DESeq2** |
| *Insl3* | x | X | x |
| *Ffar4* | x | X | x |
| *Slc45a3* | x | X | x |
| *Slc38a5* | x | X | x |
| *Itgal* | x | X | x |
| *Trf* | x | X | x |
| Slit1 | x | X |  |
| Cpz |  | X | x |
| Tgfb3 |  | X | x |
| Ces1g |  | X |  |
| Ankrd2 |  | X | x |
| Nvl |  | X |  |
| Ogdhl |  | X | x |
| Pfkfb4 |  | X |  |
| Slc33a1 |  | X |  |
| Anxa9 |  | X | x |
| Ddb2 |  | X |  |
| St3gal1 |  | X |  |
| Acsn5 |  | X | x |
| Cyp17a1 |  | X |  |
| Olfml2b |  | X | x |
| Pf4 |  | X |  |
| Nptx2 |  | X | x |
| Dnah6 |  | X |  |
| Sbpl |  | X | x |
| Adcy6 |  | X |  |
| Gm5424 |  | X | x |
| Mbp |  | X |  |
| Fbxo2 |  | X |  |
| Mycl |  | X |  |
| Eci1 |  | X |  |
| Capn12 |  | X |  |
| Col6a1 |  | X |  |
| Gpr55 |  | X |  |
| **HIGH: DRY** |  |  |  |
| **Gene ID** | **DGE edgeR** | **DTE edgeR** | **DGE DESeq2** |
| *Rin2* | x | X | x |
| *Igfbp3* | x | X | x |
| *Ctgf* | x | X | x |
| Cyp2e1 | x |  |  |
| Fmo2 | x |  |  |
| Tnfrsf21 | x |  |  |
| Cyp2f2 | x |  |  |
| Dennd2d | x |  |  |
| Nedd1 |  | X |  |
| Wdr83 |  | X |  |
| Gpx4 |  | X |  |
| Asah1 |  | X |  |
| Adm |  | X |  |
| Tmem108 |  | X | x |
| Nkx3-1 |  | X |  |
| Ybx1 |  | X |  |
| Arfgef2 |  | X |  |

**Supplemental Figure 1:**
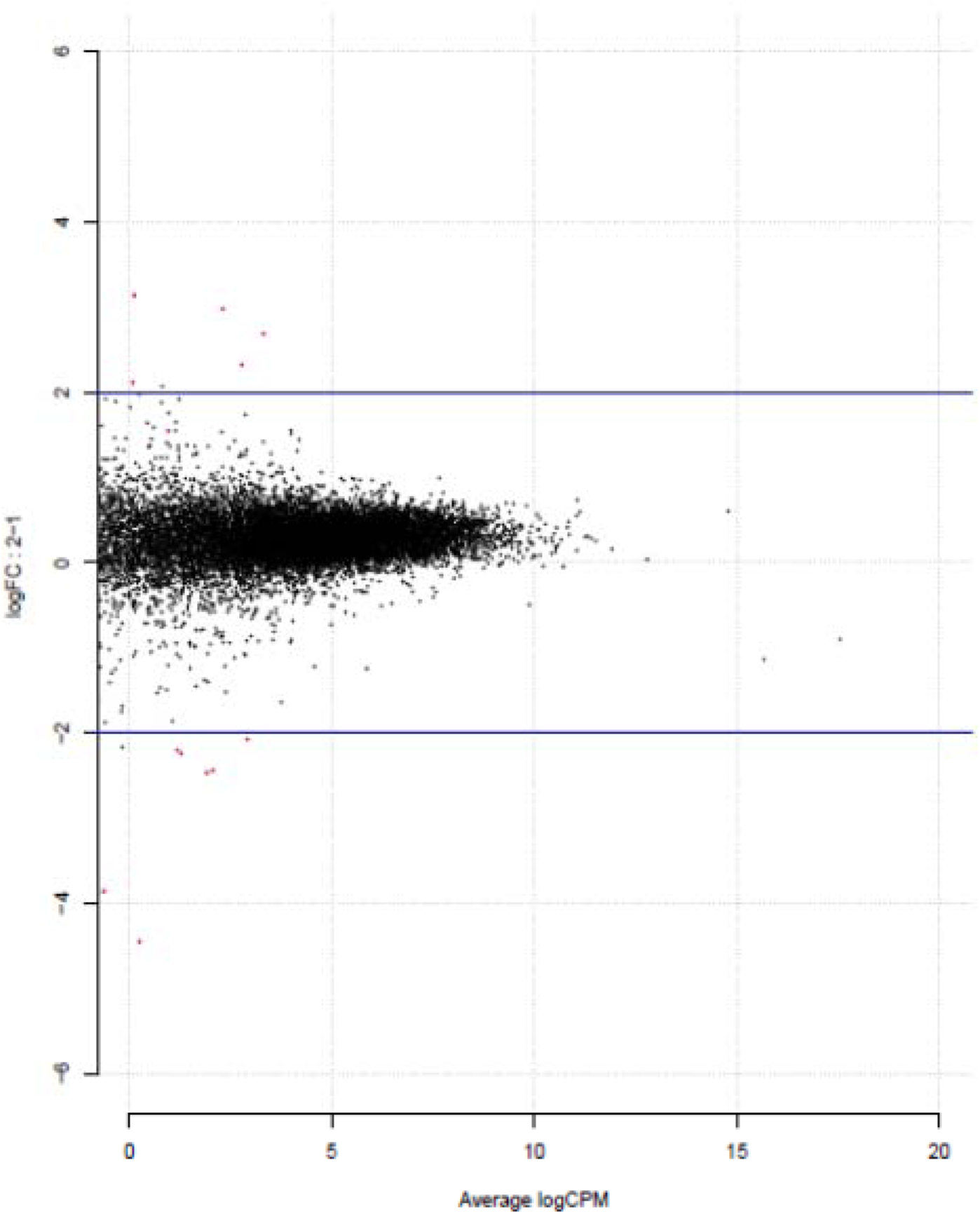
Plot of edgeR determined differentially expressed genes. The 15 significant genes are in red, with positive values indicating increased expression in the DRY group, and negative values depicting increased expression in the WET group.

**Supplemental Figure 2:**
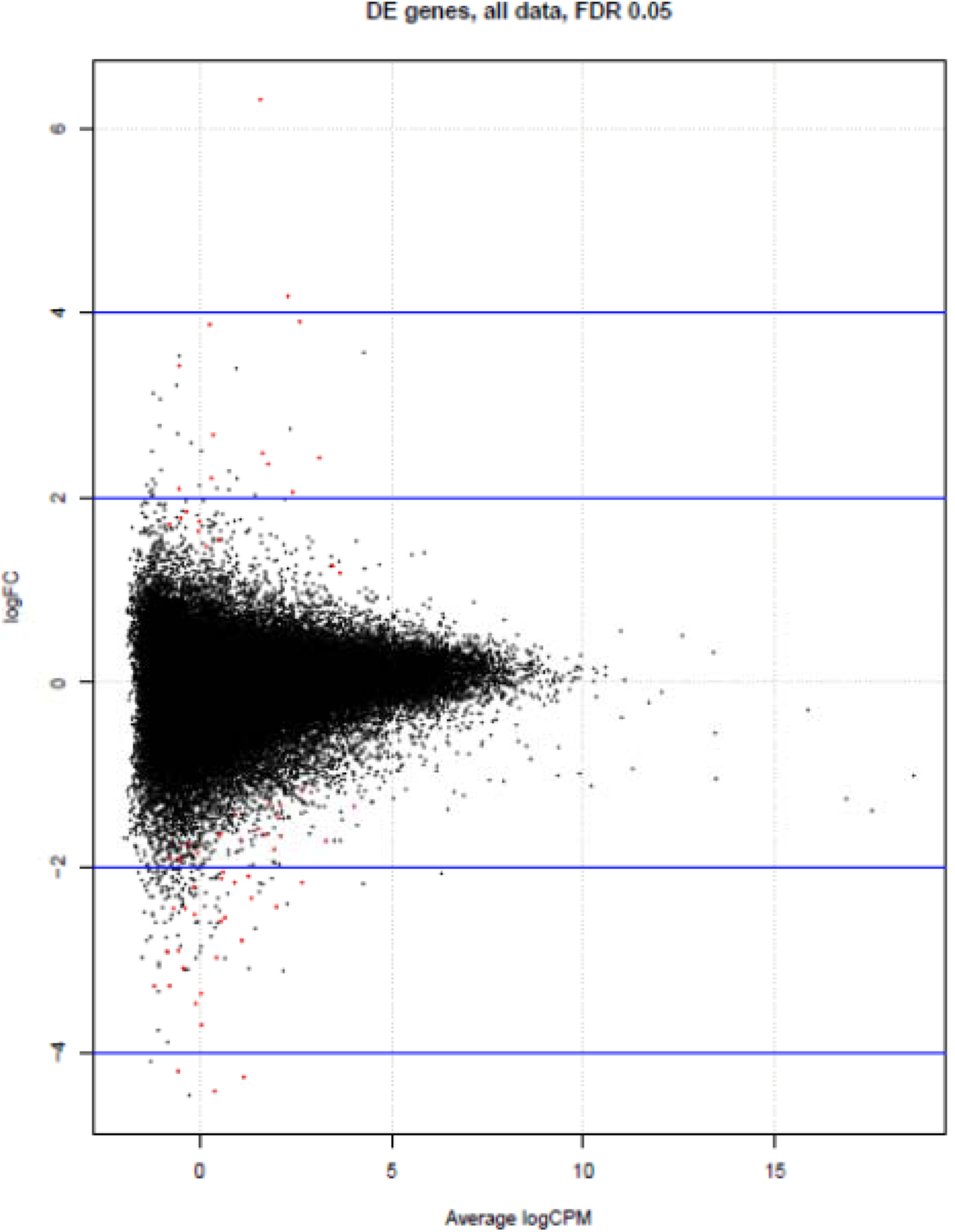
Plot of edgeR determined differentially expressed transcripts. The 66 significant transcripts are in red, with positive values indicating increased expression in the DRY group, and negative values depicting increased expression in the WET group.

**Supplemental Figure 3:**
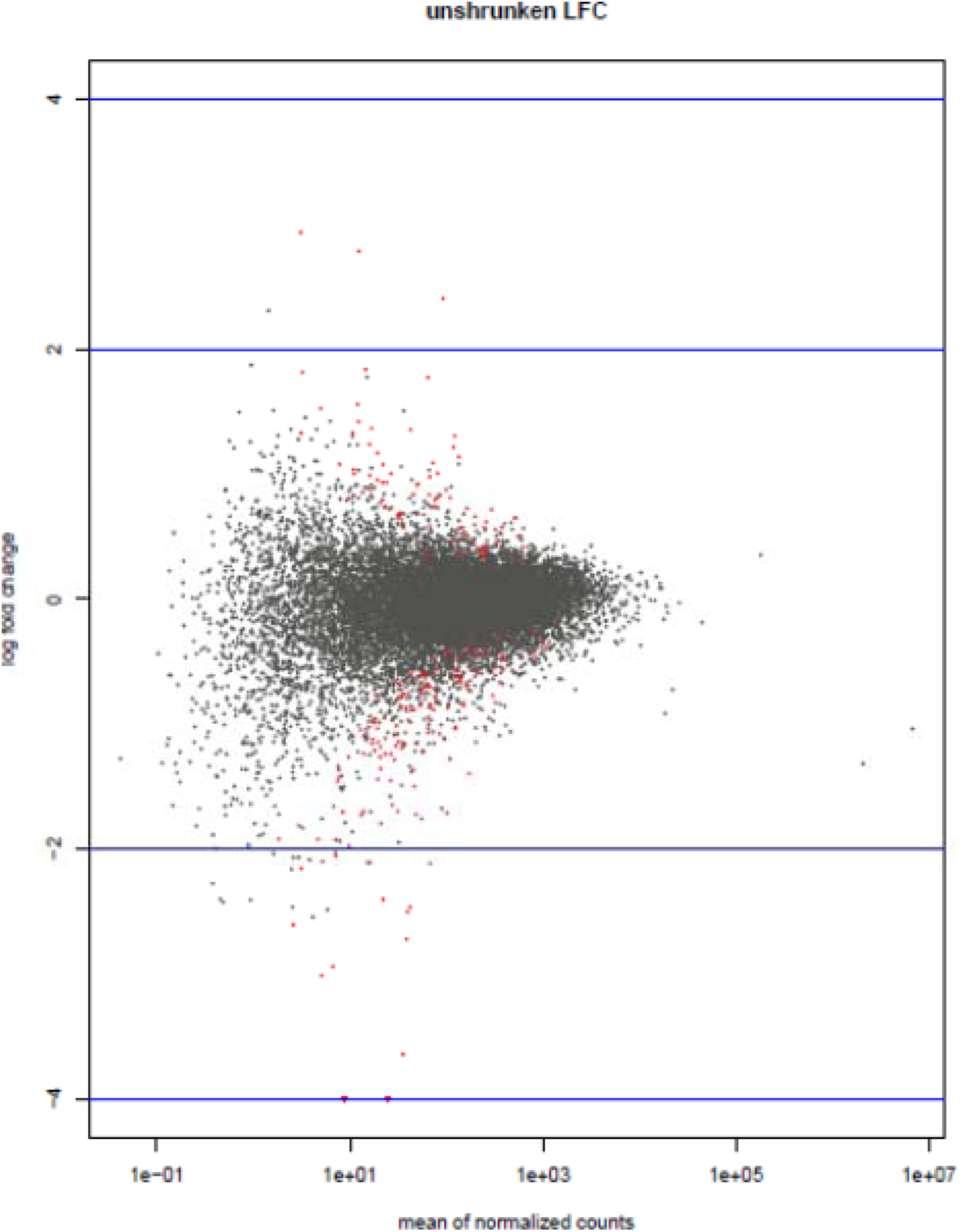
Plot of DESeq2 determined differentially expressed transcripts. The 215 significant transcripts are in red, with positive values indicating increased expression in the DRY group, and negative values depicting increased expression in the WET group.

## Supplemental DropBox Files (will be submitted to Dryad upon acceptance)

Optimized final un-annotated transcriptome (good.BINPACKER.cdhit.fasta)

Annotated transcriptome (good.BINPACKER.cdhit.fasta.dammit.fasta)

Dammit gff3 file of annotation (good.BINPACKER.cdhit.fasta.dammit.gff3)

Salmon folder including salmon quant outputs for 22 individuals (salmon)

Salmon merged quant file (NEWmergedcounts.txt)

Gene ID by Transcript ID matrix (NEWESTfinalMUS.txt)

Transcripts without matches from edgeR DTE analysis (DTEno-matchBLASTnSequences.md)

